# Abl2 mediates microtubule nucleation and repair via tubulin co-condensation

**DOI:** 10.1101/2022.06.21.496973

**Authors:** Wanqing Lyu, Daisy Duan, Kuanlin Wu, Chunxiang Wu, Yong Xiong, Anthony J. Koleske

**Author notes:** These authors contributed equally to this work (Wanqing Lyu, Daisy Duan).

## Abstract

Abl family kinases are evolutionarily conserved regulators of cell migration and morphogenesis. Genetic experiments in *Drosophila* suggest that Abl family kinases interact functionally with microtubules to regulate axon guidance and neuronal morphogenesis. Vertebrate Abl2 binds to microtubules and promotes their plus-end elongation both in vitro and in cells, but the molecular mechanisms by which Abl2 regulates microtubule (MT) dynamics were unclear. We report here that Abl2 regulates MT assembly via condensation and direct interactions with both the MT lattice and tubulin dimers. We find that Abl2 promotes MT nucleation, which is further facilitated by the ability of the Abl2 C-terminal half to undergo phase separation and form co-condensates with tubulin. A naturally occurring tubulin binding-deficient Abl2 splice isoform fails to promote nucleation. Abl2 binds to regions adjacent to MT damage and facilitates their repair via fresh tubulin recruitment and increases MT rescue frequency and lifetime. MT recovery after nocodazole treatment is greatly slowed in Abl2 knockout COS-7 cells compared to wild type cells. We propose a model in which Abl2 locally concentrates tubulin and recruits it to MT tips or to defects in the MT lattice to promote MT repair, rescue, and nucleation.

## Introduction

Microtubules (MT) are essential components of the eukaryotic cytoskeleton that play crucial roles in intracellular trafficking, cellular morphogenesis, cell migration, and cell division^1-4^. MTs are mostly nucleated from and anchored via their minus ends to the MT organizing center (MTOC), a specialized structure responsible for MT nucleation and assembly during interphase and spindle formation during mitosis^5-9^. Liquid-liquid phase separation can concentrate MT effector proteins to drive local MT nucleation^10-13^. The plus ends of MTs undergo constant turnover, termed dynamic instability to stochastically grow and shrink; switch from growth to shrinkage (catastrophe); and switch from shrinkage to growth (rescue)^2,14^. A collection of microtubule-binding proteins (MBPs) regulate MT plus end dynamics, enabling them to sample the cellular environment, deliver materials, and regulate signaling pathways^14-19^. MTs can harbor sites of lattice defects induced by mechanical stress, severing enzymes, or fast polymerization^20-24^. Tubulin can be recruited and incorporated at these sites of lattice defects and this rescue is stimulated by MBPs such as CLASP2^25-27^.

Abl family nonreceptor tyrosine kinases, Abl1 and Abl2 in vertebrates, play essential roles in the development and function of the heart, vasculature, brain, and immune system^28-34^. Adhesion and growth factor receptors signal through Abl1 and Abl2 to phosphorylate key mediator proteins that together coordinate changes in cytoskeletal structure and adhesion dynamics^35-46^. In addition to targeting these substrates, Abl2 directly binds actin and synergizes with the actin-binding protein cortactin to stabilize actin filaments and maintain dendritic spine stability^47-51^. However, the ability to regulate adhesion and actin dynamics does not fully explain the roles that Abl2 plays in neuronal development^50,52,53^. Genetic studies in *Drosophila* suggest that functional interactions between *Abl* and *orbit*, the fly ortholog of CLASP2, are required for proper axonal growth cone guidance throughout development, which requires the formation of the polarized MT network^54,55^. In the context of mitotic spindle regulation, CLASP2 enhances MT nucleation and lattice repair by recruiting tubulin to damaged sites^25,26^. Previous work from our lab showed that Abl2 directly binds to MTs and promotes MT elongation both in vitro and in cells^56,57^. These findings collectively led us to explore whether and how Abl2 impacts MT nucleation and repair.

We provide evidence here that direct binding of the Abl2 C-terminal half to MTs and tubulin dimers promotes nucleation and repair. In addition to binding the MT lattice, Abl2 binds with high affinity to tubulin dimers. We demonstrate that Abl2 forms condensates in a concentration- and salt-dependent manner, which can recruit tubulin and facilitate MT nucleation. A naturally occurring splice isoform^58^, Abl2Δ688-790, which lacks part of the high-affinity tubulin-binding region, retains binding to the MT lattice but does not bind tubulin nor promote MT nucleation. Abl2 recognizes damaged MT segments, mediates lattice repair, and increases the rescue frequency of dynamic MTs in vitro. Consistent with these findings, loss of Abl2 in cells greatly slows the recovery of MT growth after nocodazole-induced destabilization. Collectively, our data suggest that Abl2 promotes MT nucleation, repair, and rescue through the ability to bind and concentrate tubulin dimers via phase separation and promote their assembly.

## Results

### Abl2 binds to free tubulin dimers via the C-terminal half

Previous work from our lab showed that Abl2 promotes MT plus-end elongation rates in vitro and in cells, and that the C-terminal half, 557-C, is both necessary and sufficient for this activity^56,57^. To test whether Abl2 binds tubulin dimers in vitro, purified 557-C and porcine tubulin dimers were applied, separately or after mixing, to a Superdex 200 Increase size exclusion column, and their elution profiles were monitored. Purified 557-C (68 kDa) eluted with an estimated Stokes radius of 68 Å, corresponding to a globular protein of 350 kDa (**Figure 1A, S1A**). Size exclusion chromatography coupled with multi-angle light scattering (SEC-MALS) analysis revealed an estimated molecular weight of 73 kDa for the peak, corresponding to the monomeric form of 557-C (**Figure S1C**). Similarly, the peak of Abl2 was analyzed as a monomer of ∼127 kDa with an estimated Stokes radius of 78 Å (480 kDa for globular protein) (**Figure S1B**). These data indicate that Abl2 exists as an extended monomer in solution and that the C-terminal half is a chief contributor to the extended conformation. Tubulin dimers (105 kDa) eluted with an estimated Stokes radius of 42 Å, corresponding to a globular protein at about 108 kDa. Upon mixing, 557-C and tubulin were observed to co-elute in a new peak with a larger Stokes radius of 101 Å, consistent complex formation (**Figure 1A**). SDS-PAGE analysis revealed that tubulin co-eluted with 557-C in earlier fractions (**Figure 1B**).

**Figure 1.**
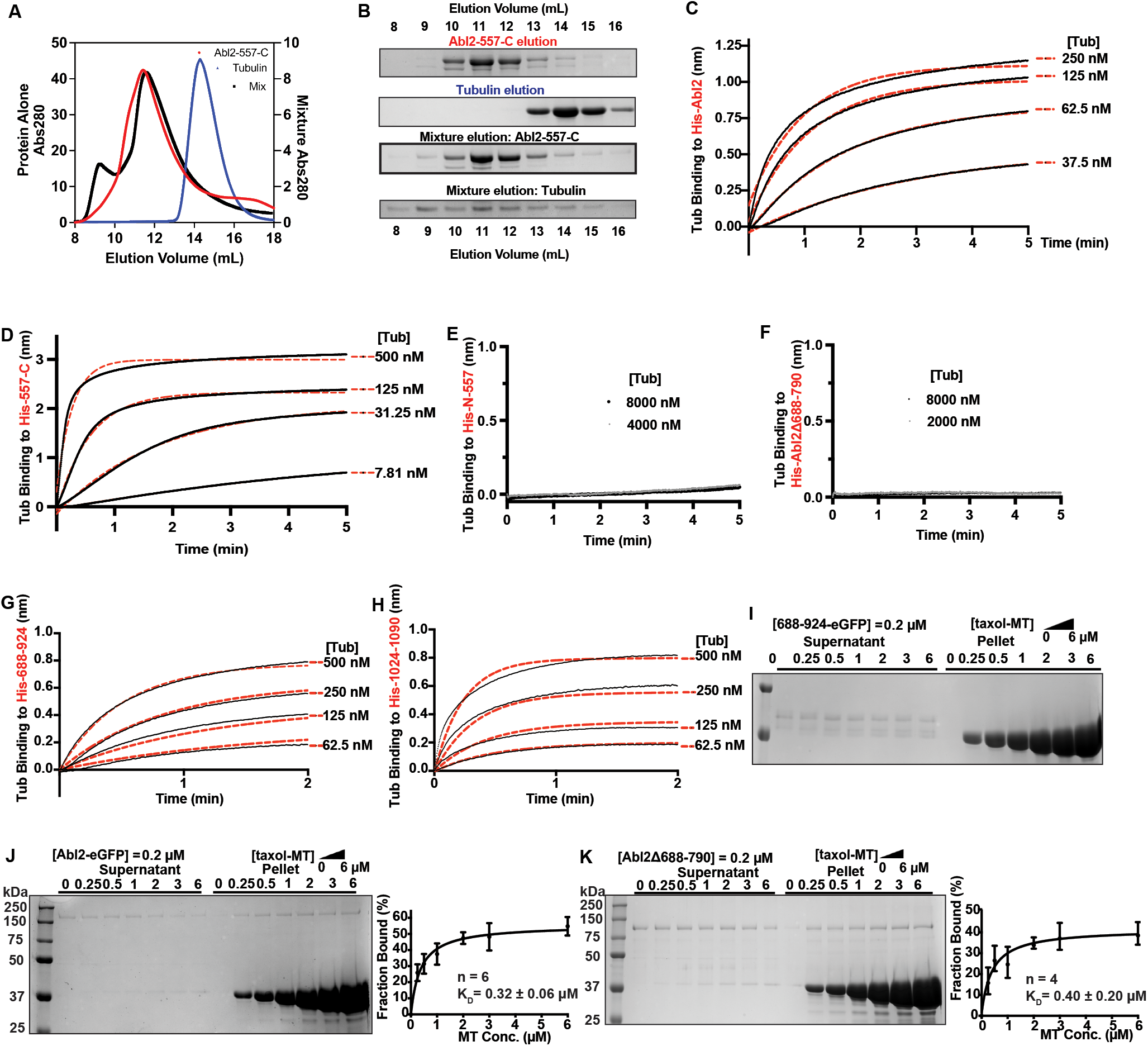
Abl2 binds tubulin via two regions in the C-terminal half. **(A)** SEC analysis of 557-C alone (left Y axis), tubulin alone (left Y axis), and the mixture (right Y axis) with **(B)** showing the corresponding SDS-PAGE analysis of the elution fractions. Tubulin alone was eluted at fractions 13-15 ml and was shifted to fractions 8-15 ml in the presence of 557-C. 6XHis-Abl2 **(C)**, 6XHis-557-C **(D)**, 6XHis-N-557 **(E)**, 6XHis-Abl2Δ688-790 **(F)**, 6XHis-688-924 **(G)**, or 6XHis-1024-1090 **(H)** were immobilized on a Ni-NTA biosensor and the association of different concentrations of tubulin was measured. Representative traces are shown, with data shown in black and global exponential fits in red dotted lines. Full concentration series (4 tubulin concentrations) were performed three independent times and used to calculate K_D_, as summarized in Table I. **(I-K)** Representative gels of the MT cosedimentation assays performed with 688-924-eGFP **(I)**, Abl2-eGFP **(J)**, and Abl2Δ688-790-eGFP in **(K)** are shown. 688-924-eGFP does not appreciably bind to MTs. Plots of percentages of Abl2-eGFP or Abl2Δ688-790 bound versus MT concentration are shown in the bottom panels with best fit K_D_ values shown. n indicates the number of experimental replicates.

We next used biolayer interferometry to measure the Abl2 binding affinity for tubulin dimers. Tubulin bound to biosensor-immobilized 6XHis-Abl2 with a K_D_ = 42 ± 13 nM. 6XHis-557-C was sufficient for high-affinity tubulin dimer binding, K_D_ = 17 ± 8 nM (**Figure 1C, D**), while the Abl2 N-terminal half, 6XHis-N-557, did not detectably bind dimers (**Figure 1E**). Two distinct Abl2 fragments, comprised of amino acids 688-924 (site I) and 1024-1090 (site II), bind independently to tubulin dimers, but with significantly reduced affinity (K_D, His-688-924_ = 118 ± 39 nM; K_D, His-1024-1090_ = 252 ± 123 nM; **Figure 1G, S1F; Table I**) relative to 6XHis-Abl2 or 6XHis-557-C. Strikingly, the naturally occurring splice isoform His-Abl2Δ688-790 binds to MTs with a similar affinity as Abl2-eGFP (K_D, Abl2-eGFP_ = 0.32 ± 0.06 µM; K_D, Abl2Δ688-790_ = 0.40 ± 0.20 µM; **Figure 1I, J; Table II**), but did not bind detectably to tubulin. On the other hand, the tubulin binding fragment 688-924-eGFP did not bind to MTs (**Figure 1H**), indicating that the MT- and tubulin-binding regions of Abl2 do not overlap completely.

**Table I.**
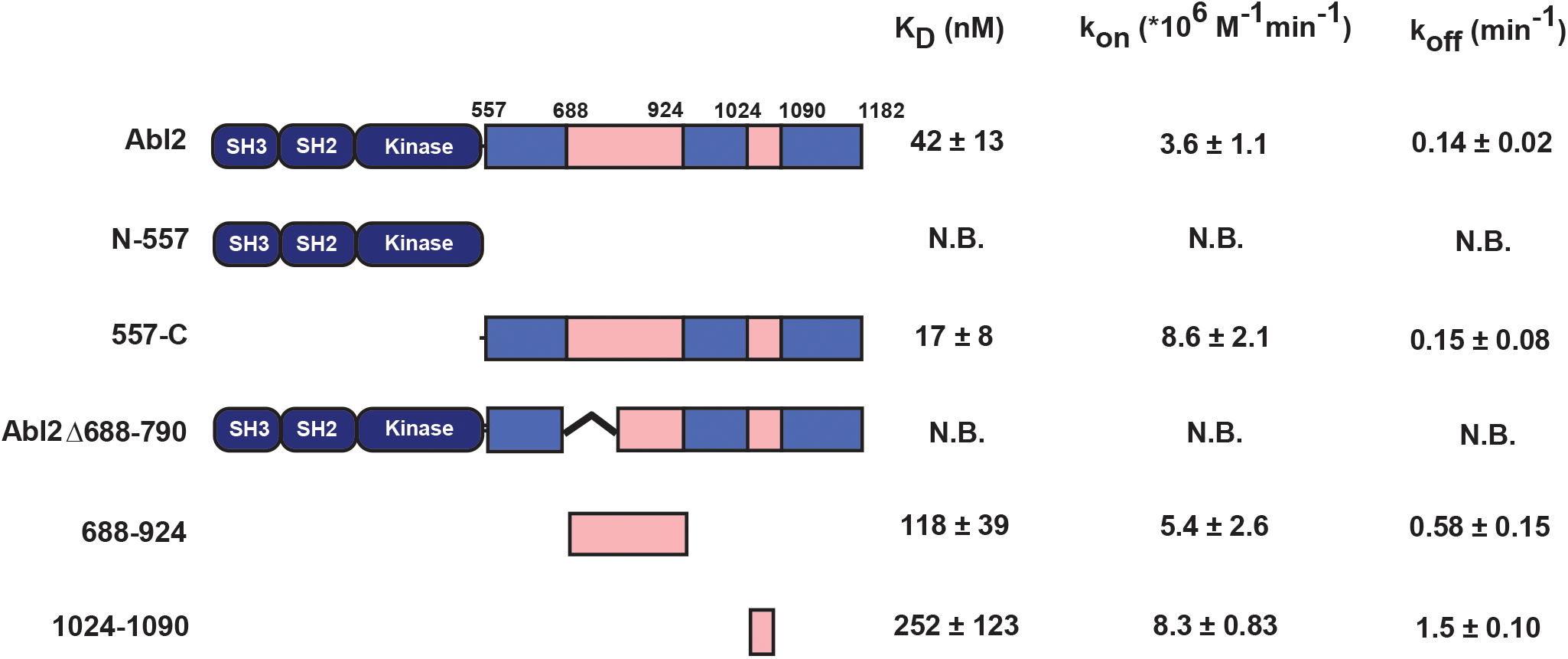
Summary of Abl2 and Abl2 fragments binding to tubulin. Disassociation constant (K_D_) data are measured as means ± S.D., for each assay condition, n = 3. N.B.: Not Binding, the binding signals are not observable or significantly low that cannot be fit.

| | | $K_D$ (nM) | $k_{on}$ ($\cdot 10^6 M^{-1} min^{-1}$) | $k_{off}$ ($min^{-1}$) |
| --- | --- | --- | --- | --- |
| Abl2 | | $42 \pm 13$ | $3.6 \pm 1.1$ | $0.14 \pm 0.02$ |
| N-557 |  | N.B. | N.B. | N.B. |
| 557-C | | $17 \pm 8$ | $8.6 \pm 2.1$ | $0.15 \pm 0.08$ |
| Abl2Δ688-790 | | N.B. | N.B. | N.B. |
| 688-924 | | $118 \pm 39$ | $5.4 \pm 2.6$ | $0.58 \pm 0.15$ |
| 1024-1090 | | $252 \pm 123$ | $8.3 \pm 0.83$ | $1.5 \pm 0.10$ |

**Table II.**
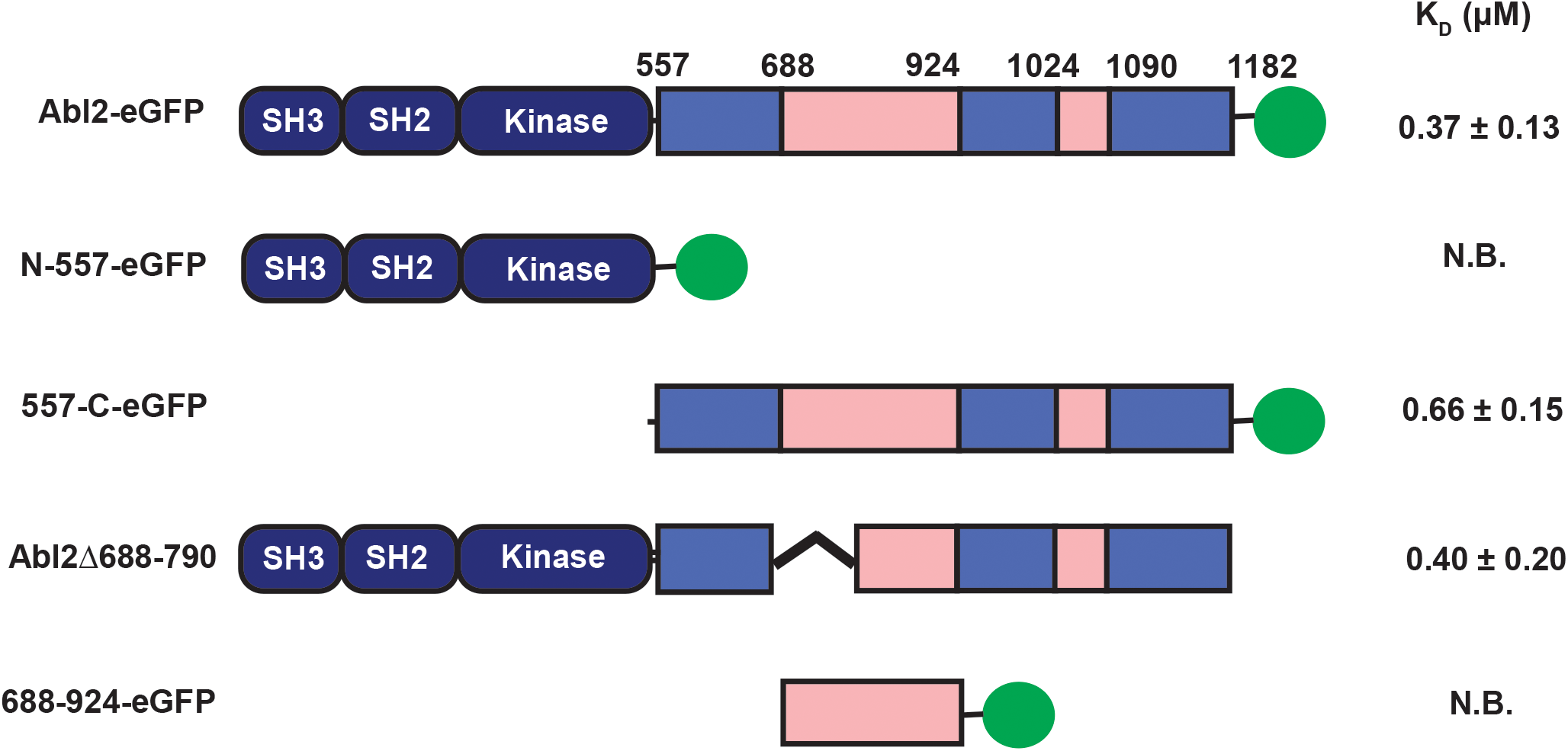
Summary of Abl2 and Abl2 fragments binding to taxol-MTs. Disassociation constant (K_D_) data are measured as means ± S.D., for each assay condition, n > 3. NB: Not Binding.

### Abl2 undergoes phase separation and co-condenses with tubulin

A growing list of MT-binding proteins regulate MT dynamics through liquid-liquid phase separation (LLPS), which is commonly associated with regions of high disorder in the protein sequence^11,59-61^. The Abl2 C-terminal half is predicted to be highly disordered using software PONDR-FIT^62^ and Diso-Pred3^63^ (**Figure 2A**). Indeed, circular dichroism (CD) spectroscopy revealed that 557-C exhibited features consistent with disordered proteins, with the commonly used CD spectra deconvolution algorithms CONTIN^64^, SELCON3^65^, and CDSSTR^66,67^ predicting 26%, 30% and 18% of disordered secondary structure content, respectively (**Figure 2B**).

**Figure 2.**
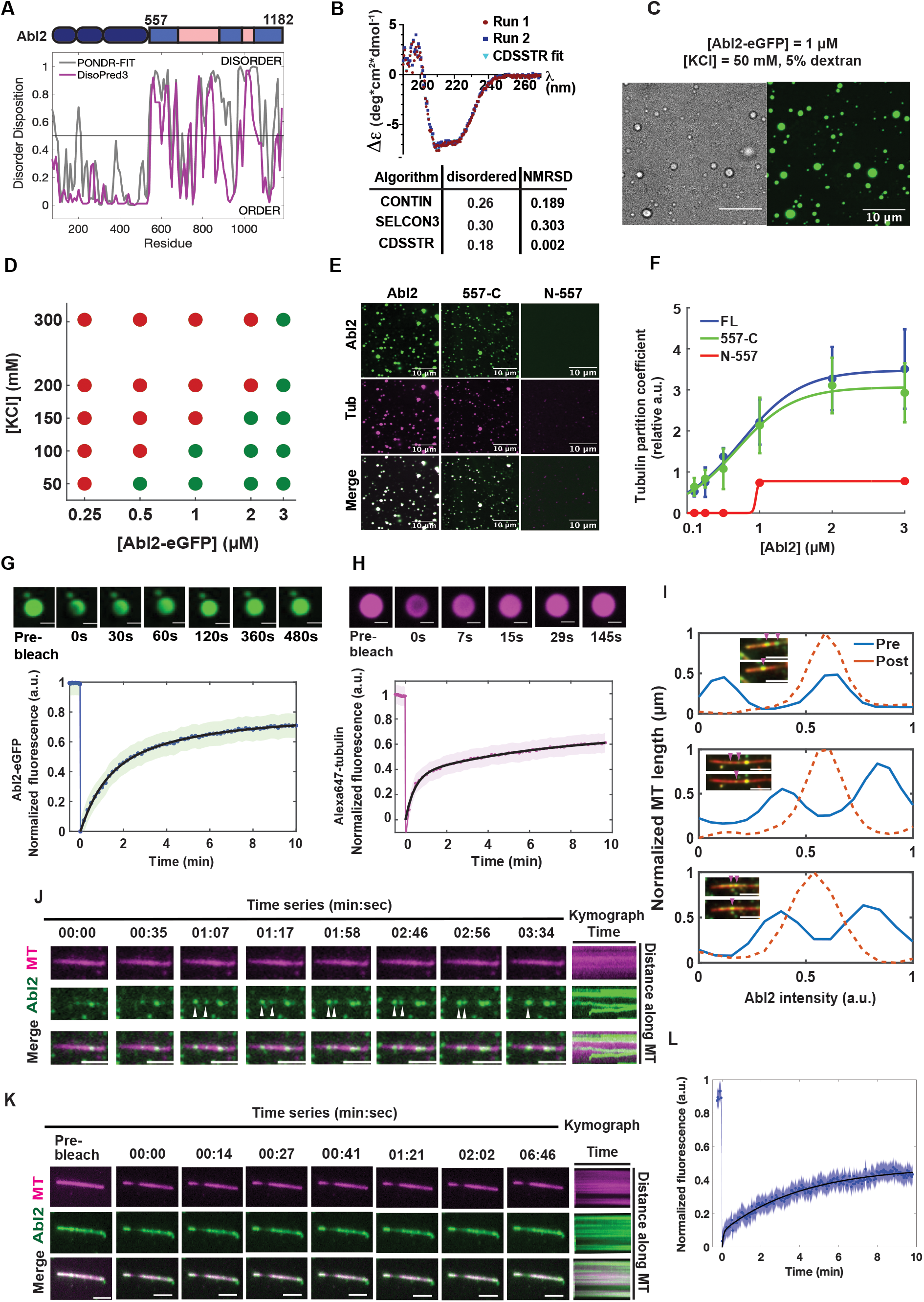
Abl2 undergoes phase separation and co-condenses with tubulin. **(A)** Disorder prediction from primary sequence of murine Abl2 using PONDR-FIT and DisoPred3 algorithms. Disorder disposition at 0.5 is used as the threshold. **(B)** Buffer-subtracted CD spectra of 6XHis-tag free 557-C collected at 4°C. CD spectra deconvolution algorithms CONTIN, SELCON3, CDSSTR analyses reveal the disordered content of the 557-C. n = 2. (**C**) 1 µM Abl2-eGFP forms condensates in BRB80, 5% dextran, 50 mM KCl, and BRB80. Condensates were imaged under brightfield and fluorescence. Scale bar, 10 µm. **(D)** Phase separation diagram of the [Abl2-eGFP] vs. [KCl] relationship. Partition coefficient (PC) 4 is defined as phase separated, shown as green dots, PC < 4 is defined as not phase separated, shown as red dots. 17-230 condensates were analyzed per condition. **(E)** Co-condensation of 1 µM Abl2, 557-C, and N-557 with equimolar concentrations of Alexa647-tubulin in BRB80, 5% dextran, 50 mM KCl. **(F)** PC analysis of co-condensed tubulin in Abl2 condensates at increasing equimolar concentrations of Abl2-eGFP and KCl. At least 90 tubulin co-condensates were scored for Abl2 and 557-C. For N557: n_2µM_ = 37; n_3µM_ = 89. Sigmoidal fits shown in solid lines. Mean ± SD shown as error bars. **(G, H)** FRAP recovery of Abl2-eGFP in condensates **(G)** and tubulin in Abl2-eGFP:Alexa647-tubulin co-condensates **(H)** in solution. Double-exponential fit shown in solid black with SEM shown as green. n 11 condensates. Scale bar, 2 µm in **(G)**; 1 µm in **(H). (I)** Representative examples of Abl2 condensate fusion events on biotinylated GMPCPP-stabilized rhodamine MTs. Condensates are indicated in magenta arrows. Post-fusion condensates yield higher mean fluorescence intensities (dashed orange line) relative to their pre-fusion condensates of various sizes (solid blue lines). **(J)** Representative time-series of two Abl2 condensates undergoing fusion on a rhodamine GMPCPP-MT (pseudo-colored magenta). Kymographs are shown on right. Scale bar, 3 µm. **(K, L)** FRAP recovery curve reveals that Abl2 condensates are undergoing dynamic exchange with other molecules on the MT and/or from solution. Double-exponential fit in solid black with SEM shown as blue. Scale bar, 3 µm. n = 15 filaments.

To investigate whether Abl2 could undergo phase separation, we observed solutions containing Abl2-eGFP under conditions believed to mimic a crowded intracellular environment (5% dextran)^68^. Abl2-eGFP puncta retained a spherical morphology as visualized in both brightfield and fluorescence imaging (**Figure 2C**). We measured the partition coefficient (PC) of Abl2 condensates, a ratio of the mean condensate intensity relative to that of background, with PC ≥ 4 as a cutoff for phase separation^11^. Using PC analysis, we observed an increased propensity to phase separate with increasing Abl2-eGFP concentration, but this was attenuated by increasing salt, a known disruptor of phase separation^68-70^ (**Figure 2D**). Fluorescence recovery after photobleaching (FRAP) of a single confocal plane within Abl2-eGFP condensate demonstrated recovery of Abl2-eGFP fluorescence with an approximate half-life t_1/2_ = 1.7 min (**Figure 2G**). When associated with GMPCPP-stabilized biotinylated MTs, Abl2-eGFP condensates moved and underwent fusion, a characteristic of phase-separated condensates (**Figure 2I, J**). The mean intensity of the larger condensate formed by the fusion of two smaller ones was greater than either alone, with a relative increase in intensity proportional to the mean intensities of pre-fusion condensates (**Figure 2I**). FRAP analysis confirmed the diffusive behavior of Abl2-eGFP on MTs, as freely exchanged with Abl2 molecules in solution (**Figure 2K, L**). Collectively, our data suggest that MTs serve as scaffolds for Abl2 to localize and condense on to possibly exert its function on MT dynamics.

Given that Abl2-eGFP binds tubulin dimers, we tested whether Abl2-eGFP could form coacervates with tubulin. When incubated under conditions that promote Abl2-eGFP phase separation (low salt, 5% dextran), Alexa647-tubulin dimers partitioned into Abl2-eGFP condensates (**Figure 2E**). The Abl2 C-terminal half is necessary and sufficient for coacervation with tubulin, as 557-C-eGFP both underwent phase separation and recruited tubulin dimers into the condensate, while N-557-eGFP did not undergo phase separation (**Figure 2D, E**). FRAP analysis of Abl2:tubulin coacervates revealed that tubulin dimers could diffuse into an internally bleached region (**Figure 2H**).

### Abl2 promotes nucleation

The finding that Abl2 has interfaces that bind tubulin dimers prompted us to investigate whether Abl2 impacts MT nucleation. We measured changes in turbidity (OD_350_), which increases upon MT polymerization, as a function of time, in the absence or presence of Abl2 (**Figure 3A**). Measurements of the critical concentration and the lag time, when turbidity reaches one-tenth of its maximum, provide sensitive measures of polymer nucleation efficiency^71-73^. Consistent with previous observations^56^, the inclusion of 1 µM Abl2 increased the plateau of turbidity, with a greater amount of polymerized MT than in the control. We confirmed that the turbidity correlated with greater MT polymerization by sedimenting the reactions and analyzing polymerized product via SDS-PAGE (**Figure S3A**). The plateau in turbidity varied as a linear function of initial tubulin concentration, yielding critical concentrations (x intercept) of 5.5 ± 1.1 µM and 0.5 ± 1.8 µM in the absence and presence of Abl2, respectively (**Figure 3B**). The lag time also decreased from 16 ± 2 min to 9 ± 3 min with Abl2 in the reactions (**Figure 3C**). Abl2Δ688-790 - which does not bind tubulin - did not promote MT nucleation over control reactions containing tubulin alone and yielded a lag time of 16 ± 2 min. The impact of Abl2 on MT nucleation was also confirmed by visualizing a portion of the reaction products via negative-stain electron microscopy (EM) at the initial phase of the reaction. Significantly more MTs were assembled when Abl2 was included in the polymerization reaction (**Figure 3D**).

**Figure 3.**
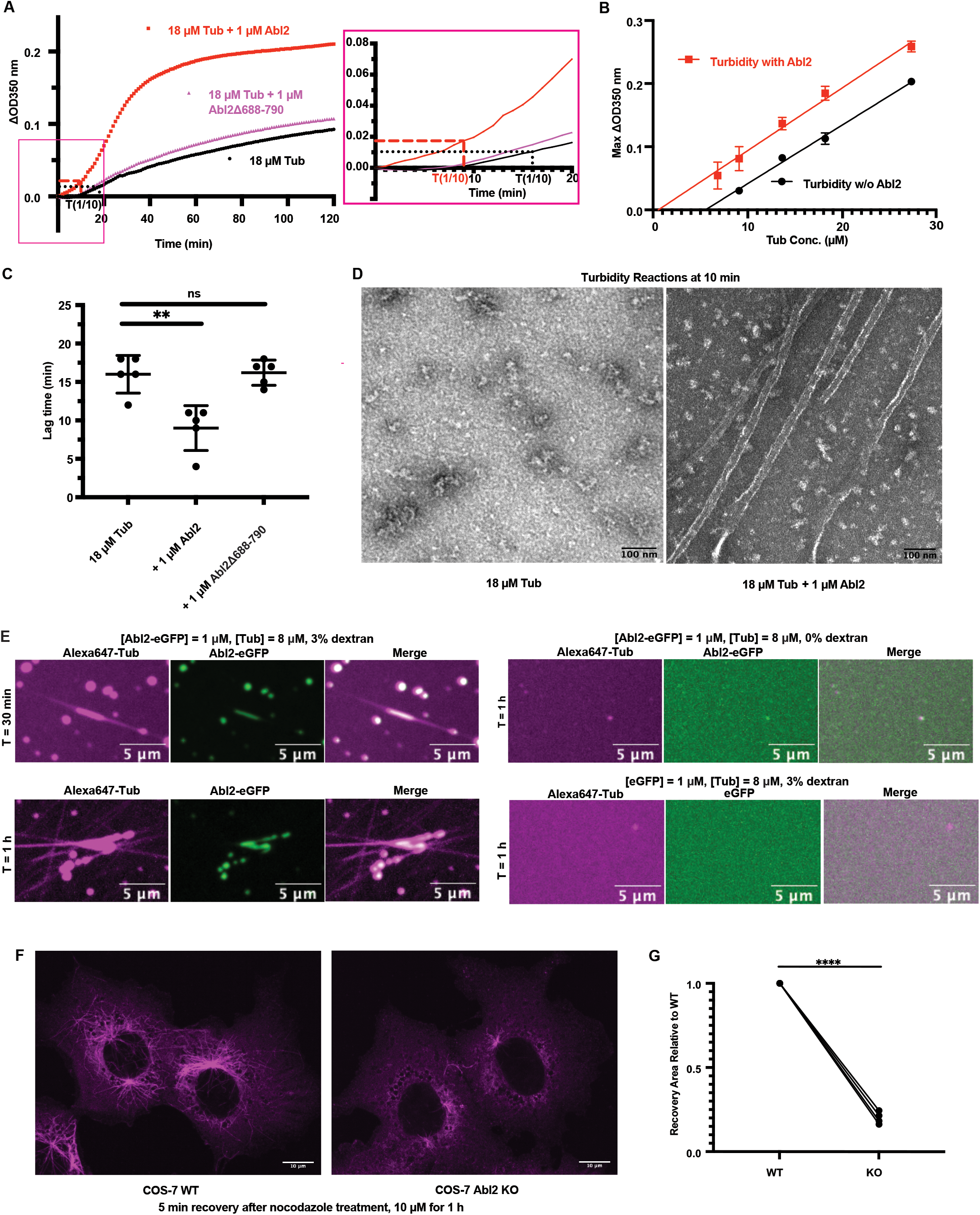
Abl2 promotes MT nucleation via interactions with tubulin and MTs. **(A)** MT assembly was monitored by measuring the increase in turbidity (ΔOD_350_). 18 µM tubulin and 2 mM GTP were incubated alone or with 1 µM Abl2 or Abl2Δ688-790. Representative time series of OD_350_ measurements are shown. Curves in the blue dotted window were are expanded in the right panel. Lag time of reactions with (red) and without Abl2 (black) are indicated with the dotted line. **(B)** Turbidity assays with different tubulin concentrations were performed and the maximal ΔOD_350_ was plotted against initial tubulin concentration to determine the critical concentration of tubulin polymerization. n = 3 replicates at each concentration. **(C)** The lag time until ΔOD_350_ reaches 1/10 of the maximal ΔOD_350_ was measured. The inclusion of Abl2 significantly decreases the lag time for MT nucleation. n = 5. **(D)** Representative samples taken 10 min after turbidity reactions were initiated were visualized under negative-stain EM. More polymerized MT segments were observed in the presence of Abl2. **(E)** Representative confocal images of the MT nucleation from the Abl2-eGFP:Alexa647-tubulin co-condensates under 3% dextran. Control reactions containing Abl2-eGFP and Alexa647-tubulin without dextran, or mixing eGFP and Alexa647-tubulin with 3% dextran did not show observable MTs growing after 1 hr. **(F)** WT COS-7 and Abl2 KO COS-7 cells were treated with 10 µM nocodazole for 1 hr, at which time nocodazole was removed and replaced with the complete medium. Immunofluorescence images of the WT and KO cells showed that 5 min after washout, MT reassembly from the microtubule organizing center was greatly reduced in Abl2 KO cells compared to WT cells. **(G)** The area of the recovered MTs in Abl2 KO cells was analyzed as outlined in Materials and Methods and Figure S2 and normalized to WT cells, by a reviewer blinded to sample identity. The experiments were repeated 4 times and each paired line represents WT/KO pair done in parallel. 10-20 cells were analyzed in each paired group. ****, p < 0.0001.

To test if co-condensation of Abl2 and tubulin facilitates MT nucleation, we measured non-templated MT nucleation using Alexa647-tubulin and Abl2-eGFP under conditions of molecular crowding (3% dextran) (**Figure 3E**). MT segments were observed growing from the Abl2:tubulin co-condensates, with numerous MTs observed after 1 h. As a control, eGFP does not form condensates under 3% dextran, and no MTs were observed after 1 h. In the absence of 3% dextran, Abl2-eGFP was diffuse in solution and few MTs were observed after a 1 hr reaction. Our results indicate that Abl2 promotes MT nucleation by interacting with tubulin in vitro, which can be further accelerated by the co-condensation of Abl2 and tubulin. We then examined how Abl2 function impacts MT reassembly following nocodazole treatment in cells (**Figure 3F**). Cells were treated with 10 µM nocodazole for 1 h and washed with the complete medium to let the MT network recover. We found that the MT recovery area in WT COS-7 cells is significantly larger than Abl2 KO COS-7 cells with comparable average cell area (**Figure 3G, S2B-D**), indicating that Abl2 promotes MT reassembly in cells.

### Abl2 mediates repair in damaged MT lattices

Given its ability to interact with both lattice and dimers, we next asked if Abl2 impacts the repair of structurally compromised MT lattices^23^. MT lattice defects were induced by incubating rhodamine-labeled MTs overnight with taxol at various temperatures and visualized as regions of reduced rhodamine intensity along the MT. We monitored repair via the incorporation of new Alexa647-tubulin at damaged sites, tracking both the number of incorporation events and the fraction of total MT length labeled with Alexa647-tubulin, termed the reporter fraction (RF). Previous reports found greater damage to the MTs stored at 23°C versus 37°C ^23,74^. In agreement with this, we observed a higher RF in the shafts of MTs stored at 23°C overnight as compared to those stored at 37°C overnight (RF_23°C_ = 0.17 ± 0.08; RF_37°C_ = 0.12 ± 0.11; **Figure S3B, C**). When mixing with Abl2-eGFP, we found that Abl2-eGFP bound at a higher density onto the more damaged 23°C-stored MTs than 37°C-stored MTs, with mean fluorescence intensities of 0.91 ± 0.03 a.u.*µm^-1^ and 0.71 ± 0.03 a.u.*µm^-1^, respectively (**Figure S3C**).

Abl2-eGFP did not uniformly decorate the damaged MT segments. To analyze this distribution, we characterized the MT shafts based on healthy and damaged regions and measured whether Abl2 has a preference. We made segmented line scans of MTs and defined segments of mean fluorescence intensities equal to or above 0.5 as ‘healthy’ and those below as ‘damaged’. ‘Borders’ are defined as segments directly adjacent to damaged ones, whose lengths are one-fifth of the total healthy segment length away from damaged sites, with ‘Lattices’ are defined as the remaining three-fifths middle portion of the healthy segment (**Figure 4A**). Abl2-eGFP bound preferentially to borders, with mean intensities of 0.39 ± 0.02 a.u. and 0.38 ± 0.02 a.u. on 23°C- and 37°C-stored MT damaged borders, respectively; and mean intensities of 0.30 ± 0.03 a.u. and 0.28 ± 0.03 a.u. on 23°C- and 37°C-stored MT lattices, respectively (mean ± SEM, **Figure 4B**). These data suggested that Abl2 detected borders of MT defects. We next asked whether Abl2-eGFP can mediate tubulin incorporation to these damage sites, so we incubated 23°C- and 37°C-stored damaged MTs with and without 1 µM Abl2-eGFP (**Figure 4C**). We discovered that the fraction of total MTs containing Alexa647-tubulin increased 1.27- and 1.26-fold to RF_23°C, Abl2_ = 0.22 ± 0.07 and RF_37°C, Abl2_ = 0.15 ± 0.07, respectively (**Figure 4D, S3D**); with the frequency of new tubulin incorporation sites into existing shafts increased in the 37°C storage condition, from f_incorp, tub, 37°C_ = 0.19 ± 0.04 events*µm^-1^ of MT length to f_incorp, Abl2, 37°C_ = 0.23 ± 0.03 events*µm^-1^ (**Figure 4E, S3E**), suggesting that Abl2 promotes repair of structurally compromised MT lattices.

**Figure 4.**
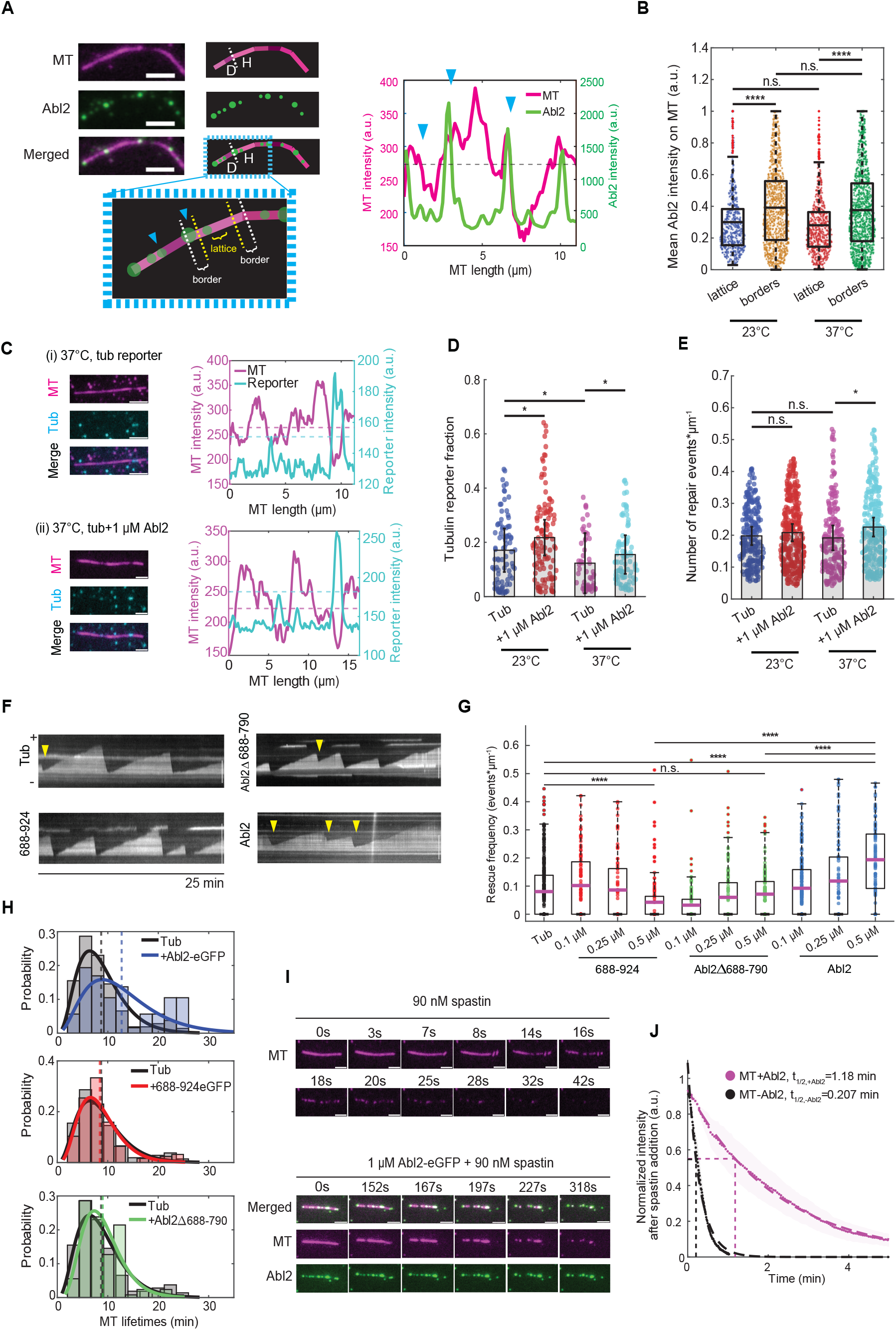
Abl2 promotes damaged lattice repair and increases MT lifetime. **(A)** Representative Abl2-eGFP localization on a 37°C-overnight stored taxol-stabilized MT. Borders are defined as boundaries on ‘healthy’ (“H”) structurally intact MT segments adjacent to ‘damaged’ (“D”) segments, demarcated with blue arrows. Lattices are stretches that are at least 1/5 of the total MT segment length away from either terminus. Scale bar, 3 µm. **(B)** Mean intensities of 1 µM Abl2-eGFP at the borders and lattices of healthy MT segments along filaments stored at 23°C and 37°C overnight. Means are shown as solid horizontal black lines, 25-75% quartiles are shown as box plots. n 600 healthy segments were analyzed per condition. Wilcoxon rank sum test performed. ****, p < 0.0001. **(C)** Representative taxol-stabilized rhodamine-MTs stored at 37°C overnight (magenta) and the repaired tubulin reporter (cyan). MT mixture was supplemented with 2 µM Alexa647-tubulin (cyan), 10 mM GTP, and 10 µM taxol, and were allowed to incorporate at damaged sites for 3 hr at 37°C with **(ii)** or without **(i)** 1 µM Abl2-eGFP. Fluorescence intensities of the MT and 647-tubulin were plotted. Dashed magenta line denotes the MT intensity threshold to distinguish healthy from damaged segments. Dashed cyan line denotes the reporter intensity threshold to score whether 647-tubulin reporter is present or not. Scale bar, 3 µm. **(D)** Total tubulin reporter fraction per MT was quantified per storage condition in the presence or absence of 1 µM Abl2-eGFP. Mean ± SEMs shown as error bars. n 50 filaments were analyzed per condition. Wilcoxon rank sum test performed. *, p < 0.05. **(E)** Total number of repair/incorporation events per MT was quantified per storage condition in the presence or absence of 1 µM Abl2-eGFP. n 200 incorporation events were analyzed per condition. Mean ± SEMs are shown as error bars. Wilcoxon rank sum test performed. *, p < 0.05; n.s., no significance. **(F)** Representative kymographs of dynamic MT filaments in presence of 10.5 µM rhodamine tubulin alone, or with 0.5 µM 688-924-eGFP, Abl2Δ688-790, or Abl2-eGFP. Scale bar, 8 µm. **(G)** Rescue frequencies (f_res_) were quantified for 10.5 µM rhodamine tubulin alone, or supplemented with 0.1, 0.25, and 0.5 µM 688-924-eGFP, 0.5 µM Abl2Δ688-790, or 0.5 µM Abl2-eGFP. While the f_res_ decreased with increasing concentrations of tubulin-binding fragment 688-924-eGFP and was not impacted under the presence of tubulin-binding deficient Abl2Δ688-790, 0.5 µM Abl2-eGFP increased f_res_ 2-fold. n 75 filaments analyzed per condition. Means shown as magenta lines, 25-75% quartile shown as box plots. Wilcoxon rank sum test performed. ****, p < 0.0001. **(H)** MT lifetime distributions of filaments grown in presence of 0.5 µM Abl2-eGFP, 0.5 µM 688-924-eGFP, and 0.5 µM Abl2Δ688-790 relative to tubulin alone. Gamma distributions were fit to MT lifetime histograms, shown in solid blue, orange, and green curves respectively, relative to tubulin control (solid black). Mean MT lifetimes are denoted as dashed lines. **(I)** 90 mM spastin supplemented with 2 mM ATP was added into TIRF chambers containing biotinylated, rhodamine GMPCPP-stabilized MTs (pseudo-colored magenta) with or without 1 µM Abl2-eGFP. For severing assays that included Abl2, Abl2 condensates were allowed to incubate for 5 min prior to imaging. Representative time series of MTs are shown. Intensity decay curves of severed MTs quantified in (J). Scale bar, 3 µm. **(J)** Mean intensity decays of MTs in absence and presence of 1 µM Abl2-eGFP shown in black and magenta scatter dots, and SEMs shown as black and magenta bands, respectively. Single exponential decay curves were fit to the data, shown as dashed black and magenta lines. n 200 filaments were analyzed per condition.

### Abl2 increases rescue frequency to prolong MT lifetimes

Repair of the damaged MT leads to the formation of GTP-tubulin enriched islands inside MT segments, which function as the GTP-caps to combat the catastrophe of MTs, therefore, resulting in a higher MT rescue frequency. To further explore our findings that Abl2 supports robust MT lattice integrity, we tested how Abl2-eGFP impacts rescue frequency (f_res_) using biotinylated GMPCPP seeds and 10.5 µM rhodamine-labeled tubulin. Consistent with our prior work^56^, Abl2 decreases the MT depolymerization rate 41.2%, from 14.11 µm*min^-1^ to 8.29 µm*min^-1^ (**Figure S4A**). Interestingly, the inclusion of Abl2 increased rescue frequency in a concentration-dependent manner (**Figure 4F, G**): incubation of MTs with 0.5 µM Abl2 increased the rescue frequency. f_res_ increased 2-fold relative to tubulin alone (f_res, 0.5 µM Abl2_ = 0.192 ± 0.022 events*µm^-1^; f_res, tub_ = 0.095 ± 0.019 events*µm^-1^, **Figure 4G**). However, the inclusion of 0.5 µM of the tubulin-binding 688-924-eGFP or the MT-binding Abl2Δ688-790 did not impact rescue frequency (f_res, 0.5 µM 688-924_ = 0.086 ± 0.05 events*µm^-1^; f_res, 0.5 µM Abl2Δ688-790_ = 0.070 ± 0.017 events*µm^-1^). These data suggest that Abl2 requires both its tubulin- and MT-binding modalities for dynamic MTs to promote MT rescue events. We also measured the net effects of Abl2 and Abl2 fragments on MT lifetime distribution (MTLD). MTs alone have a mean lifetime of 8.7 min. The addition of 0.5 µM Abl2 significantly extended the MTLD, a mean increase of 46.7% to 12.7 min, indicating that MTs grown in the presence of Abl2 are older. In contrast, the inclusion of 0.5 µM Abl2-688-924-eGFP did not alter MTLD, while the addition of 0.5 µM Abl2Δ688-790 increased the mean lifetime slightly to 9 min (**Figure 4H**).

Considering its ability to maintain MT lattice integrity, we wondered whether Abl2 could protect against active MT lattice damage mediated by the ATP-dependent MT severing enzyme spastin. We first incubated biotinylated GMPCPP-stabilized rhodamine MTs alone or in the presence of 1 µM Abl2-eGFP prior to flowing in 90 nM spastin and 2 mM ATP and tracked the decay in MT fluorescence intensity over time to monitor the impact of spastin on MT disassembly (**Figure 4I**). Spastin promoted rapid disassembly of MTs with t_1/2_ =0.21 min, while pre-incubation of MTs with 1 µM Abl2-eGFP greatly slowed the spastin-mediated disassembly rate to t_1/2, Abl2_=1.18 min (**Figure 4J**). This observation suggests that Abl2 delays spastin-mediated severing, suggesting that Abl2 can protect MT lattices from active damage.

## Discussion

We demonstrate here for the first time that Abl2 undergoes phase separation and forms coacervates with tubulin to promote MT nucleation. In addition to this role in nucleation, Abl2 mediates lattice repair, and its impact on both functions likely explains its ability to promote MT rescue frequency. Our results are consistent with a model in which Abl2 promotes the recruitment and addition of tubulin to MTs in different functional scenarios.

### Abl2 interacts with tubulin through two distinct regions

We identified two tubulin-binding regions in Abl2, regions containing amino acids 688-924 (Site I) and 1024-1090 (Site II), each sufficient to bind tubulin dimers itself, albeit with weaker affinity than Abl2 and the C-terminal half. Curiously, the Abl2Δ688-790 splice isoform did not bind detectably to tubulin even though it retains site II (**Figure 1; Table I**). We speculate that Abl2 initially interacts with tubulin via Site I, which has a higher affinity to tubulin. The binding of tubulin to Site I may be required to expose Site II to an additional tubulin dimer and promote their interactions in a head-to-tail fashion (**Figure 5A**). This model may help to explain why the naturally occurring splice isoform Abl2Δ688-790, which lacks all or part of Site I, significantly disrupt binding to tubulin and render it unable to promote MT nucleation. The protein sequence of the two tubulin-binding sites in Abl2 showed no significant alignment with known tubulin-binding domains, indicating novel interactions between tubulin and tubulin-binding proteins.

**Figure 5.**
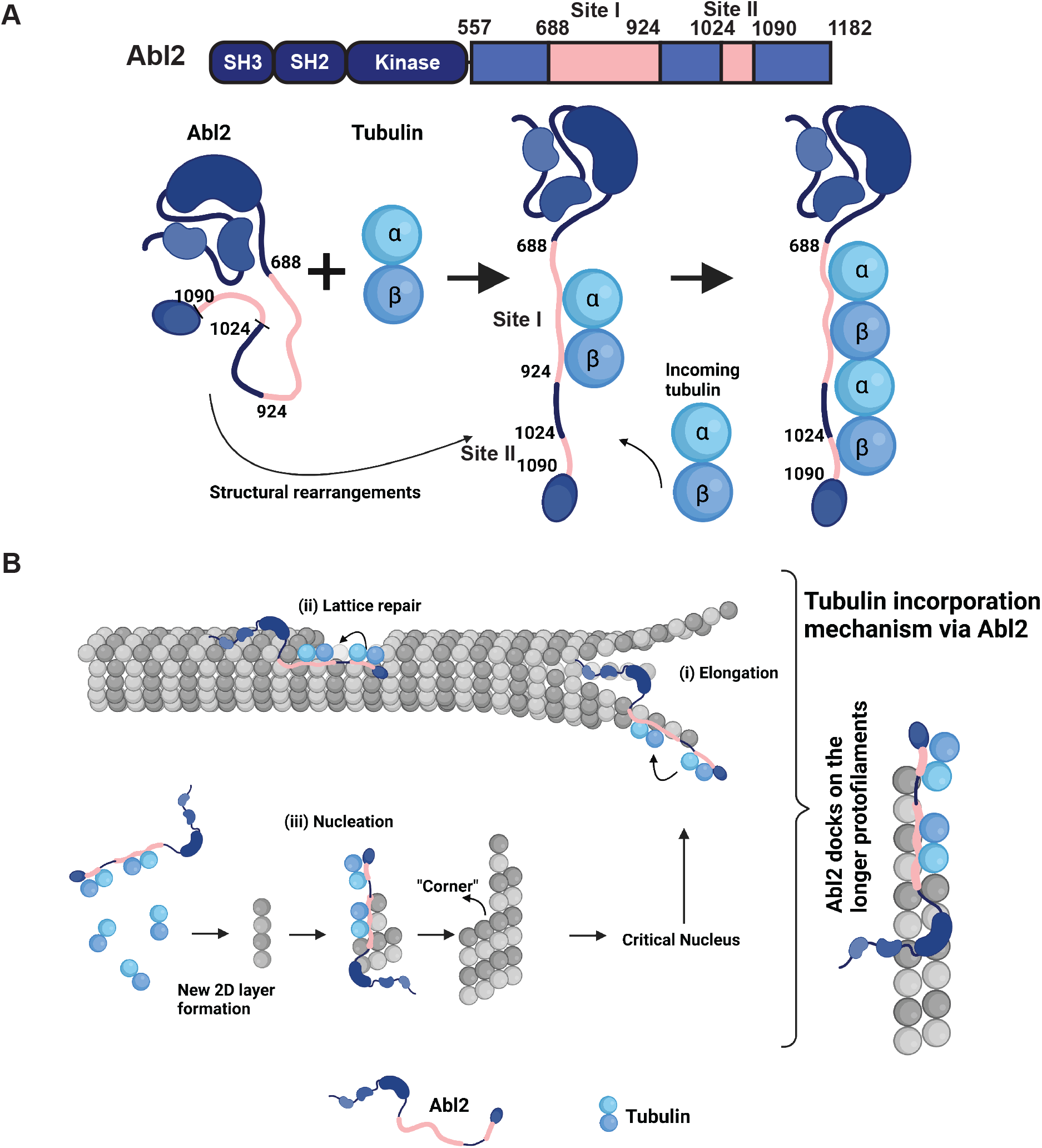
Model for Abl2 in regulating MT dynamics. **(A)** Abl2 directly interacts with tubulin dimers using its solvent-exposed 688-924 (site I). The 1024-1090 region (site II) is less accessible in the native conformation. Binding to a single tubulin dimer via site I may induce a conformational change within 557-C, leading to solvent exposure of site II, which can further recruit an additional dimer. **(B)** Abl2 can dock onto the longer protofilaments **(i)** on growing MT ends; **(ii)** MT lattice defects borders; and **(iii)** the tubulin oligomers to facilitate the incorporation of new tubulin dimers for MT elongation, lattice damage repair, and nucleation.

### Phase separation is critical to Abl2 function in regulating MT dynamics

Under conditions of molecular crowding, we found that Abl2 undergoes salt- and concentration-dependent phase separation and can recruit tubulin into the dense phase. Protein condensation is often driven by intrinsically disordered regions and multivalent binding regions^61,68-70,75^. Consistent with this, we found that the Abl2 C-terminal half is significantly disordered and has two distinct sites for tubulin binding (**Figure 2**). Phase separation appears to be a common feature of diverse MT regulators including TPX2^18,76^, tau^59,60,77^, and CLIP-170^78^, with diverse activities including MT nucleation, protection of MT severing, and condensing tubulin dimers for MT growth. We showed that new MTs nucleated from Abl2-eGFP:tubulin co-condensates. Under the same conditions, eGFP alone was not able to phase separate or interact with tubulin to promote MT assembly (**Figure 3E**). We also showed that Abl2 condensates readily associated with MTs (**Figure 2I-L**). Protection of MTs from spastin damage after incubating 1 µM Abl2-eGFP suggests that phase-separated Abl2 clusters act as steric blockers to prevent spastin localization and severing (**Figure 4I, J**). Phase separation could potentially serve as a potential regulatory mechanism for Abl2 to locally modulate MT dynamics.

### Does phase separation impact Abl2 kinase signaling?

In its kinase-inactive conformation, the Abl2 N-terminal half adopts a locked conformation in which its N-terminal Src homology (SH) 3, SH2, and kinase domains engage in the intramolecular interactions that inhibit kinase activity^79-82^. Signaling through integrin and growth factor receptors is believed to relieve this inhibition and trigger extensive phosphorylation of downstream effectors^78,83-85^. It is unclear how phase separation would impact the ability of Abl2 to interact with upstream activating receptors or its substrates. It is also unknown if phase separation behavior could be modulated by signaling cues, i.e. recruitment to the plasma membrane via binding to the cytoplasmic regions of integrin or growth factor receptors or by post-translational modifications, including phosphorylation^61,68,75,86-89^. We note that the predominant tyrosine phosphorylation sites in Abl2 occur in its N-terminal half, which does not phase separate (**Figure 2E**). Database searches indicate multiple potential serine and threonine phosphorylation sites in the C-terminal half, which may impact its ability to undergo phase separation and/or bind MTs or tubulin^86,90^.

### Abl2 interacts with MTs and tubulin to promote MT nucleation, growth, and lattice repair

Our lab previously reported that Abl2 directly binds MTs and promotes MT elongation rates^56,57^. We show here that Abl2 reduces the critical concentration for spontaneous tubulin polymerization and shortens the lag time for MT nucleation. The tubulin binding-deficient Abl2Δ688-790 isoform does not impact critical concentration or lag time, indicating that Abl2 requires its ability to bind tubulin dimers to mediate robust nucleation in vitro (**Figure 3**). Classical models for MT nucleation proposed that new MTs form via a nucleation-elongation mechanism, in which the formation of a critical nucleus is the rate-limiting step. Recent studies demonstrated that ‘the critical nuclei’ are first formed as 2D layers of a growing lattice prior to its maturation into a cylindrical tube ^91,92^. Enlargement of 2D lattices is energetically favorable but kinetically impeded by the difficulty in adding new protofilament layers. We propose that Abl2 may facilitate this by binding a nascent protofilament and recruiting and condensing tubulin dimers to promote layer formation. In addition, Abl2 may then bind to the lattice and facilitate tubulin addition at corners of 2D lattices, which may be structurally akin to the corners at growing MT ends and damaged holes (**Figure 5B, iii)**. We note that growing MT ends contain a heterogeneous mixture of straight ends, curled multi-protofilament sheets, and flared or ragged ends^93,94^. The structures of these damaged sites are understudied and are likely to contain a heterogeneous mixture with corners for tubulin incorporation. We speculate that Abl2 recognizes one or more of these structures within the damage region, bringing along tubulin to mediate rapid tubulin dimer addition (**Figure 5A, i and ii**).

### Does Abl2 mediate actin-MT crosstalk?

Previous work showed that Abl2:actin interactions are critical for the formation of dynamics actin-based protrusions in diverse cell types^51,57,95^. Abl2 also synergizes with cortactin to bind and stabilize actin filaments in vitro (and interact with another actin-interactor, cortactin, to maintain dendritic spine stability^50^. Cooperative interactions of Abl2 and actin filaments are reported through two distinct regions: aa. 688-924 and 1090-1182^47,51^. Our finding that the regions in Abl2 that bind tubulin dimers overlap significantly with those that bind actin filaments raises the fundamental questions of whether and how Abl2 mediates actin:MT crosstalk or whether the two polymer systems compete for Abl2. It is also unclear how the different interactions of Abl2 with actin versus MTs contributes to its ability to regulate cells and neuronal morphology and function.

In summary, we demonstrate that Abl2 and tubulin co-condense, acting as compartmentalized reactors to: 1) nucleate MTs; and 2) promote the incorporation of tubulin dimers at damaged lattice sites for repair and rescue. Our study provides a mechanistic model to probe how Abl2 regulates MT assembly and repair and provides tools to explore how these mechanisms regulate cell morphogenesis and migration.

## Acknowledgments

This work was supported by the National Institutes of Health grants R21 NS112121, R01 MH115939, R01 NS105640, and R56MH122449 (to A.J.K.), an American Heart Association Predoctoral Fellowship (to W.L.), and a National Science Foundation Graduate Research Fellowship Program (to D.D.). We would like to thank Josie E. Bircher for providing thoughtful feedback on the manuscript, and Xianyun Ye for her assistance in purifying DNA preparations. We also greatly appreciate the Titus Boggon Lab for generously providing us access to the BLItz instrument.

## Author Contributions

W.L., D.D., A.J.K designed experiments. W.L., D.D. conducted experiments and wrote the manuscript. W.L., D.D., A.J.K. prepared all expression constructs, and W.L. and D.D. purified all protein. W.L., D.D., A.J.K. revised and edited the manuscript. C.W., K.W. prepared reagents for and assisted with SEC-MALS and fixed cell nucleation experiments and analyses, respectively.

## Materials and Methods

### Tubulin purification and labeling

Porcine brain tubulin was prepared as previously described in Castoldi & Popov (2003)^96^. 2 ml 14 mg*ml^-1^ porcine brain tubulin was labeled with TAMRA as described in Peloquin et al. (2005)^97^. Biotinylated tubulin was obtained from Cytoskeleton.

### Molecular cloning and Abl2 purification

Murine Abl2 (residues 74-1182), Abl2-eGFP, N terminal half (N-557), N-557-eGFP, C terminal half (557-C), 557-C-eGFP, and fragments contain amino acids 688-790, 688-924, 688-924-eGFP, 1024-1090 were cloned with an N-terminal 6XHis tag into the pFastBac1 vector (Invitrogen) for insect cell expression. Abl2Δ688-790 was generated using PCR-based mutagenesis and confirmed by DNA sequencing. Abl2 and Abl2Δ688-790 were cloned into pN1-EGFP expression vector for mammalian cell expression. Recombinant baculoviruses expressing Abl2 or Abl2 fragments in pFastBac were generated using the Bac-to-Bac expression system in Sf9 insect cells according to the manufacturer’s instructions (ThermoFisher, Waltham, MA). After 36-48 hr infection with baculoviruses, Hi5 cells were collected and centrifuged for 3,000 rpm for 5min at 4°C. Cells were lysed using buffer containing 20 mM HEPES pH 7.25, 5% glycerol, 500 mM KCl, 20 mM imidazole, 1 mM DTT, 1 mM PMSF, 1X protease inhibitor cocktail, 1% Triton-X100 and left to incubate at 4°C on a rotisserie stand for 5-10 min. Lysates were ultracentrifuged using Ti70.1 rotor for 45 min, 40K rpm at 4°C. After collecting and 0.45 µm filtering the supernatant, proteins were passed through disposable columns with Ni-NTA resin beads via gravity flow three times. Resin was washed in order with the following buffers: wash A (20 mM HEPES pH 7.25, 5% glycerol, 500 mM KCl, 20 mM imidazole, 1 mM DTT, 1 mM PMSF, 1X protease inhibitor cocktail); B (20 mM HEPES pH 7.25, 5% glycerol, 1 M KCl, 20 mM imidazole, 1 mM DTT, 1 mM PMSF, 1X protease inhibitor cocktail), A, and C (20 mM HEPES pH 7.25, 5% glycerol, 300 mM KCl, 20 mM imidazole, 1 mM DTT, 1 mM PMSF, 1X protease inhibitor cocktail). Bound proteins were step eluted off the Ni-NTA resin (Invitrogen) using wash C buffer containing 300 mM imidazole in 5×1 ml fractions. His-Abl2 and Abl2 fragments for biolayer interferometry assays were exchanged into storage buffer containing 20 mM HEPES pH7.25, 5% glycerol, 100 mM KCl, 1 mM DTT using either Superdex 75 increase 10/300 GL column for proteins with MW < 75kDa, or Superdex 200 increase 10/300 GL column for proteins with MW ≥ 75 kDa. His tags were removed by adding TEV protease into purified Abl2 constructs, and were incubated on a rotisserie stand at 4°C for 2.5-4 hrs. His-tag cleaved proteins were exchanged into the storage buffer containing 20 mM HEPES, 5% glycerol, 300 mM KCl, and 1 mM DTT using either Superdex 75 increase 10/300 GL column or Supderdex 200 increase 10/300 GL column. Peak fractions were collected, snap frozen, and stored at -80°C until use.

### Spastin purification

pGEX-PP-Spastin(87-616)DeltaExon4 was purchased from Addgene (item ID #128794). Spastin was transformed and expressed into *E. coli* BL21 cells. 4l expression cultures were grown in 2XYT media at 37°C, 200 rpm until A_600_ reached 0.6, which was then followed by induction with 0.5 mM IPTG. Cultures were grown 16°C for 16 hr. Cells were harvested by centrifugation at 4K rpm in SLA1500 rotor 15 min at 4°C. Cell pellets were resuspended in lysis buffer (50 mM Tris-HCl pH 8.0, 5% glycerol, 0.1% Triton X-100, 5 mM MgCl2, 1 mM DTT, 1X PMSF, 1X protease inhibitor cocktail). Lysates were clarified for 20 min in SA600 rotor at 15K rpm at 4°C, and 0.45 μm filtered. 2 ml of glutathione agarose resin was equilibrated using lysis buffer without added detergent, and pipetted into a disposable gravity flow column. Lysates were passed through column 3X. Resin was washed with buffers in the following order: A (50 mM Tris-HCl pH 8.0, 300 mM KCl, 5 mM MgCl_2_; 5% glycerol, 1X DTT, 1X PMSF); B (50 mM Tris-HCl pH 8.0, 1 M KCl, 5 mM MgCl_2_; 5% glycerol, 1X DTT, 1X PMSF); A; and C (50 mM Tris-HCl pH 8.0, 150 mM KCl, 5 mM MgCl_2_; 5% glycerol, 1X DTT, 1X PMSF). 5 mL of Wash C supplemented with 1 mg PreScission protease was added into the resin and left to incubate on the column overnight at 4°C. After ∼16 hr, GST-cleaved spastin was removed from PreScission protease by passing through Superdex 200 increase 10/300 GL column and stored in 20 mM HEPES, 5% glycerol, 300 mM KCl, 5 mM MgCl_2_, 1 mM DTT. The first three peak fractions were concentrated using 0.5 ml centrifugal filter units (EMD Millipore #UFC501096) that were pre-equilibrated with storage buffer, and stored in -80°C until use.

### TEV protease purification

The plasmid pRK793 TEV S219V was obtained as a gift from the Boggon Lab and transformed into BL21 *E. coli* for purification. The protein was expressed in 2 l of BL21 *E. coli* culture followed by 0.2 mM isopropylβ-D-thiogalactopyranoside (IPTG) induction at 37°C for 3 hrs. Cells were sonicated in buffer containing 20 mM Tris pH 8, 5 mM βME, 500 mM NaCl, 4 mM imidazole, 1 mM DTT, 1 mM PMSF, 1X protease inhibitor cocktail. Lysates were ultracentrifuged using Ti70.1 rotor for 45 min, 40K rpm at 4°C. After collecting and 0.45 µm filtering the supernatant, proteins were passed through disposable columns with Ni-NTA resin beads via gravity flow 3X. Resin was washed with the following buffers in the following order: wash A (20 mM Tris pH 8, 5 mM βME, 500 mM NaCl, 4 mM imidazole, 1 mM DTT, 1 mM PMSF), B (20 mM Tris pH 8, 5 mM βME, 250 mM NaCl, 40 mM imidazole, 1 mM DTT, 1 mM PMSF), and C (20 mM Tris pH 8, 5 mM βME, 40 mM Imidazole, 100 mM NaCl, 1 mM DTT, 1 mM PMSF). Bound proteins were eluted off the Ni-NTA resin using elution buffer (20 mM Tris pH 8, 5 mM βME, 400 mM Imidazole, 100 mM NaCl, 1 mM DTT, 1 mM PMSF, and 1X protease inhibitor cocktail). The eluted fractions were then applied to a mono S cation exchange column for clean up using buffer A (20 mM HEPES pH 7.25, 5% Glycerol, 1 mM DTT) and buffer B (20 mM HEPES pH 7.25, 5% Glycerol, 1 M KCl, 1 mM DTT). The eluted fractions containing TEV were pooled, flash-frozen in liquid nitrogen, and stored at 80°C.

### Microtubule cosedimentation assays and quantification

Cosedimentation assays were performed as previously described^57,98^. Double-cycled GMPCPP (Jena Bioscience, Thuringia, Germany) stabilized MTs were grown as described^99^. Taxol-MTs were polymerized at a final concentration of 60 µM at 37°C in polymerization buffer [80 mM PIPES, pH 6.8, 1 mM MgCl2, 1 mM EGTA, 1 mM GTP, and 15 nM paclitaxel (taxol)]. The taxol-stabilized MTs and GMPCPP-stabilized MTs were set aside for cosedimentation. For MT cosedimentation assays, 0.2 µM Abl2 or Abl2 fragments were mixed with increasing concentration of MTs (0 to 6 μM) at 37°C for 20 minutes in binding buffer [80 mM PIPES, pH 6.8, 70 mM KCl, 1 mM GTP, 5 nM taxol (100 μL reaction volume)]. Mixtures were pelleted by high-speed centrifugation at 120,000 X *g* for 20 minutes at 37°C. Pellet and supernatant fractions were recovered and separated by SDS-PAGE, stained with Coomassie Blue G-250 (Bio-Rad Laboratories, Hercules, CA) then destained in water. The SDS-PAGE gels were then scanned with Bio-Rad ChemiDoc™ Touch Imaging System and quantified by densitometry using ImageJ software. Binding affinity was quantified either as the percentage of Abl2/Abl2 fragments bound to MTs over total amount of Abl2/Abl2 fragments in the reaction for each concentration or as the amount of Abl2 bound to MTs for each concentration of Abl2. Experiments were repeated at least 4 times for each experimental condition (n ≥ 4). Equation 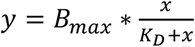 (**Equation 1**) was used to fit the curve^100^, where y is specific binding, x is the ligand concentration, B_max_ is the maximal binding (same units as y), and K_*D*_ is the binding affinity (same units as x). Binding curves, affinities (K_D_), and R^2^ values for curve fitting were calculated using GraphPad Prism 9 GraphPad.

### Tubulin binding analysis with size-exclusion chromatography

Size-exclusion chromatography used a Superdex 200 increase 10/300 GL column equilibrated in 20 mM HEPES pH 7.25, 5% glycerol, 100 mM KCl, 1 mM DTT. The column was calibrated with standard proteins of known Stokes radii (Sigma-Aldrich, St. Louis, MO). Abl2 or Abl2-557-C and tubulin were mixed with tubulin at a ratio of 1:4 and incubated for 30 min on ice, and then injected onto the column. Control experiments were performed with each protein alone. The collected fractions were analyzed by SDS-PAGE, stained with Coomassie brilliant blue G250 (Sigma-Aldrich, St. Louis, MO), and scanned.

### Tubulin binding affinity measurements using biolayer Interferometry

The biolayer interferometry technique using the BLItz system (ForteBio) was used to measure binding kinetics for the tubulin interaction with 6XHis-Abl2 and 6XHis-Abl2 fragments. His tag-binding Ni-NTA biosensors were hydrated in binding buffer (20 mM HEPES pH 7.25, 5% glycerol, 100 mM KCl, 1 mM DTT, 0.02% Tween) for 10 min. For each tubulin concentration (ranging from 7 nM to 2000 nM), the following procedure was performed. An initial baseline was collected by immersing the biosensor in binding buffer for 1 min, and then 4 µl of fixed concentrations of His_6_-Abl2 or His_6_-Abl2 fragments (0.3 µM or 1 µM) were loaded to the biosensor for 5 min. The Abl2-loaded biosensor was returned to binding buffer for collection of a second baseline for 1 min and then placed in 4 μl of tubulin for a 5-min association step. For each data point, the background binding was also measured. For each tubulin concentration, the difference in the signal (in nm) just prior to the association step and that at the end of the association step was subtracted from the difference in signal for background binding. The binding curves were plotted fit to the one-phase exponential curves assuming a shared koff using the “association kinetics (two or more ligand concentrations)” model with GraphPad Prism to obtain a dissociation constant.

### Turbidity assay and preparation of Abl2-MT sample grids for EM imaging

18 µM porcine brain tubulin in BRB80 was incubated with the MT polymerization buffer (2 mM of GTP, 1 mM DTT, 15% Glycerol) alone or in the presence of Abl2 and Abl2 fragments at 37°C. Tubulin assembly was monitored by measuring turbidity at 350 nm (A_350_) for 2 hours using SpectraMax M6 Multi-Mode Microplate Reader recording spectrophotometer. Control experiments were done by monitoring A_350_ for buffer alone with 0.5 μM Abl2 and Abl2 fragments without tubulin. 4 µl reactions at 10 min were taken out and visualized using electron microscopy (EM) of negatively stained samples. A 400-mesh copper grid (Ted Pella, Redding, CA) overlaid with a very thin continuous carbon layer was gently glow discharged and 4 µl of the diluted protein was applied to the grid. After a 30s adsorption, the reactions were blotted away from the grid with filter paper (Whatman No.1) leaving a thin layer of solution on the grid. 4 µl of 2% uranyl-acetate solution were applied to the grid for 30 s before blotting twice. After blotting, the grid was left to dry for 2 min. The negative stain sample of Abl2 was imaged using a Tecnai12 transmission electron microscope (TEM) and images recorded on a Gatan CCD camera at ∼-2-3 µm defocus.

### Construction and preparation of flow chambers for TIRF imaging

Glass microfluidic chambers were constructed as described previously in Johnson-Chavarria et al. (2011)^101^. Briefly, inlet and outlet ports were generated into polydimethylsiloxane (PDMS) molds using a blunt tip needle. Holes were drilled into a glass cover slide with a diamond tip bit. Plasma cleaning was used to bond PDMS molds onto the cover slides where PDMS ports and holes meet.

Coverslips were cleaned with the following solutions, all incubated in a sonicator for 15 min (unless otherwise stated) and washed with ddH_2_O in between steps: 2% Hellmanex, ddH_2_O, 90% ethanol, 1M HCl (1 hour to overnight). To extensively rinse off HCl, coverslips were washed with ddH_2_O 3X for 15 min per step. Coverslips were then cleaned with 0.22 µm filtered 1M KOH for 30 min, followed by ddH_2_O, and 100% ethanol for 60 min to overnight. Coverslips were dried in a 55°C incubator and silanized with 300mL dichlorodimethyl silane in 300 mL hexane solution for 35-45 min. They were subsequently sonicated in hexanes for 15 min. Coverslips were air dried and stored in sterile 50 mL Falcon tubes at -20°C until use. Immediately before use, TIRF chambers were assembled using parafilm, with rectangular cutouts to allow for flow between inlet and outlet ports; and was sealed between clean coverslips and the PDMS chamber with heat (via pressing the chamber with coverslip-side down onto surface of a 100°C heat block for ∼5-10 s).

Flow chambers were prepared by washing in BRB80 and incubating 1 mg*mL^-1^ biotin-BSA (0.22 μm filtered in BRB80) for 5-10 min. Chambers were blocked with 2% Pluronic F-127 and allowed to incubate at RT for 30-45 min, followed by washing with BRB80 and functionalizing with 50 mg*ml^-1^ neutravidin (0.22 µm filtered in BRB80). After 5 min incubation, flow chambers were washed with BRB80 and perfused with biotin GMPCPP-MT seeds. Density of seeds were checked under the scope. Chambers were washed with BRB80 to flow out unattached seeds.

### Microtubule dynamics assay

Microtubule seeds were prepared by mixing 10% biotin, 1% rhodamine-labelled 10 µM tubulin in BRB80 buffer, supplemented with 1 mM GMPCPP. The seed mixture was incubated in a 37°C water bath for 30-45 min, followed by centrifugation in a TLA100 rotor at 80K rpm for 5 min at 37°C. Single-cycled seeds were resuspended in warm BRB80 buffer and ready for use. Imaging buffer consisted of 10.5 µM 10% rhodamine-labelled tubulin, 1 mM GTP, 0.02% methylcellulose, 1X anti-blink cocktail (1% β-ME, 40 mM glucose, 250 nM glucose oxidase, 64.5 nM catalase, 1 mM Trolox, BRB80), and BRB80. All proteins were centrifuged in TLA100 rotor for 5 min at 80K rpm, 4°C prior to use to remove aggregates. In conditions with Abl2-eGFP, Abl2Δ688-790, or Abl2-688-924-eGFP, salt concentration was adjusted such that the final [KCl] in the imaging buffer was 50 mM. Reaction mixture was pipetted up and down 3X prior to perfusing into flow chamber. Dynamic MTs were allowed to grow, equilibrating in the chamber for 5 min prior to image acquisition. The objective was maintained at 37°C with an objective heater. Images were acquired at 1 fps for 25min, 2×2 binning with 300 ms integration times per channel, using the Nikon Elements software; and a Nikon Eclipse TE2000-S Inverted Microscope equipped with a 100X TIRF 1.49NA oil objective and Andor Zyla 4.2 sCMOS camera. Dynamic instability parameters were analyzed from kymographs generated from Multi Kymograph using ‘read velocities from tsp’ plug-in (tsp050706.txt macro written by J. Rietdorf and A. Seitz, EMBL). At least 3 technical replicates (at least one different TIRF chamber per independent experiment) were conducted for each condition. MT lifetime distributions were fitted to a gamma distribution.

### Preparation of damaged MTs and lattice repair assay

100 µl mixture of 2% biotinylated, 10% rhodamine-labeled 20 µM tubulin was thawed on ice and incubated with 1 mM GTP. The mixture was allowed to polymerize in 37°C water bath for 30-45 min. MTs were centrifuged in a TLA100 rotor for 5 min at 37°C and resuspended in warm BRB80 buffer supplemented with 10 µM paclitaxel (taxol). Taxol-stabilized MTs were separated into 2 mixtures: 1) one 50 µl stored at RT (23°C) overnight; 2) the remaining 50 µl stored in 37°C.

The following day, the MTs were centrifuged for 5 min at 80K rpm, 37°C; and resuspended in warm BRB80 supplemented with 10 µM taxol. The 2 mixtures were separated into 2 25 µl Eppendorf tubes. Alexa647-tubulin and Abl2-eGFP were clarified for 5 min at 80K rpm, 4°C. 2 µM Alexa647-tubulin and 10 mM GTP were each added into each 25 µl mixture of MTs. 1 µM Abl2-eGFP was added into one of each 23°C and 37°C-stored MT reactions (except for damaged boundary analysis assays, where 0.5 µM of Abl2-eGFP was added in absence of Alexa647-tubulin). All reactions were left to incubate in a 37°C water bath for 3 hours prior to imaging. TIRF chambers were functionalized as described above. 25 µl of each reaction (control and +Abl2 conditions) containing MTs stored at either 23°C or 37°C were perfused into the chamber. Warm BRB80 buffer was flowed into the chamber to wash out unattached taxol-MTs. Still images for the green (Abl2), red (MT; pseudo-colored magenta in), and far red (tubulin reporter; pseudo-colored cyan) channels were acquired using single-band emission filter sets for 488, 561, and 638 nm, with 2×2 binning 300 ms acquisition times on the Nikon Eclipse TE2000-S Inverted microscope.

To score for damaged MTs at either temperature, 3-pixel wide line scans across the entire length of rhodamine-MTs and tubulin reporter were recorded. User-defined MT and tubulin reporter intensity thresholds were used as parameters in custom-written MATLAB script to determine borders of MT segments that are structurally damaged, and whether tubulin reporter was present or not. MT segments of mean fluorescence intensity lower than user-defined MT-intensity threshold were scored as ‘damaged’. If the mean intensity of tubulin reporter of the equivalent damaged MT length is at least the user-defined reporter intensity threshold, the reporter length, reporter fraction, and frequency of reporter incorporation for that one MT were recorded.

For Abl2 localization and border analysis, MT segments of mean fluorescence intensity at least the user-defined MT threshold value were determined as ‘healthy’. Along these ‘healthy’ segments, borders are defined as edges of healthy segments that border damaged segments (of mean intensity values less than user-defined threshold). Border length was quantified as being 1/5 of the total healthy MT segment away from either terminus, with the lattice length quantified as the middle 3/5 of the MT segment. Localization of 0.5 µM Abl2-eGFP was quantified by taking the mean fluorescence intensity of the green channel along sub-sections of the healthy MT segment that were either borders or lattices.

### Microtubule severing assay

10% biotinylated, 10% rhodamine labelled GMPCPP-stabilized MTs were flowed into TIRF chambers, followed by flow-in and incubation of 1 µM Abl2-eGFP in imaging buffer (1X anti-blink cocktail, 0.02% MC, 50 mM KCl, and BRB80) for 5 min to allow for condensate formation. Flow chamber was washed with BRB80 buffer to remove non-bound Abl2. 90 nM spastin in imaging buffer supplemented with 2 mM ATP was introduced into the chamber and image acquisition at 0.34 fps for 15 min immediately followed. 3 pixel-wide line scans of MTs were recorded for MTs at the beginning of each image acquisition and fluorescence intensity decay traces were plotted and fit to a single exponential decay: 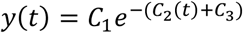 (**Equation 2**).

### Phase separation assay

Abl2-eGFP, 557-C-eGFP, and N-557-eGFP were thawed on ice and concentrated to final concentration of 10 µM or higher using 0.5 ml centrifugal filter units (EMD Millipore #UFC501096) equilibrated with Abl2 storage buffer (20 mM HEPES, 300 mM KCl, 5% glycerol, 1 mM DTT). Concentrated proteins were centrifuged to remove aggregates in a TLA100 rotor at 80K rpm, 4°C for 5 min. Individual wells in Cellvis #1.5 glass-bottom 96-well plates were washed with BRB80 and blocked with 2% Pluronic F-127 for 30-45 min. 20 µl of imaging buffer (0.2% methylcellulose, 1X anti-blink cocktail, BRB80, 0.1-3 µM Abl2-eGFP, 557-C-eGFP or N-557-eGFP) was mixed on ice. 6.25 µl 16% Dextran-70 was added into the imaging buffer reaction to achieve final concentration of 5% dextran. The mixture was pipetted up and down 3X at RT and added into the well. Condensates were left to age for 10 min at RT prior to image acquisition. Epifluorescent imaging was performed using the Nikon Eclipse TE2000-S Inverted microscope adapted to a mercury lamp source and a 50:50 beam splitter to toggle between TIRF and Epi modalities. Focus was set just above the coverslip at the bottom of the well (z-height ≈ 3795-3850mm) where condensates have sedimented and settled.

To calculate partition coefficients and score for phase-separated condensation, we thresholded still images of sedimented Abl2 coacervates by filtering out particles for sizes 1-00 and those with circularity values of 0.1-100. We then analyzed particles using Otsu, black and white thresholding, followed by color inversion. Circular ROIs were generated to quantify background values. Mean intensities of detected particles and background were calculated for each condition. If mean(particles) ≥ mean(background), *i*.*e*. PC ≥ 4, the particle was scored to be phase-separated condensate under the particular Abl2 concentration and salt concentration condition. For Abl2-tubulin co-condensation analysis, all partitioning experiments were maintained at 50 mM KCl, 5% dextran. Tubulin PCs were fitted to a sigmoidal equation, 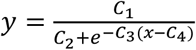 (**Equation 3**).

### Circular dichroism of Abl2 557-C

Purified His-tag free Abl2 557-C was purified into storage buffer containing 20 mM HEPES pH 7.25, 300 mM KCl, 5% glycerol, 1 mM DTT. CD spectra were collected at a constant temperature of 4°C using a Chirascan CD spectrometer (Applied Photophysics) at 0.5 nm wavelength step size and 0.5 s per wavelength point. CD spectra of the gel filtration buffer were collected to background subtract from the sample spectra. Averages of 2 spectra from 280 to 190 nm were calculated per replicate for duplicate technical replicates.

### FRAP of Abl2 and tubulin in solution

Images were acquired using a Cellvis #1.5 glass-bottom 96-well plate. Condensates were prepared and aged as described above, maintained at 50 mM KCl. Prior to photobleaching, images were acquired every second for 30 seconds to obtain an average pre-bleach fluorescence intensity value. Photobleaching was carried out using 20% power of the 405 nm laser line, with 20 μs dwell time at a single focal plane near top of the condensate. An ROI was chosen as a circular region within the condensate. Intensity of each ROI was recorded every 10s (∼0.14 fps) over a 10-15 min acquisition period. Recovery in photobleached ROIs was normalized to changes in background intensity to account for global bleaching. Images were acquired using Nikon SoRA spinning disk confocal microscope equipped 100X oil objective. Triplicate technical replicates of Abl2 and duplicate replicates of tubulin FRAP curves were fitted to double exponential decay: 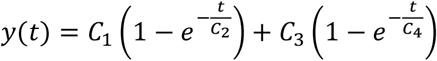 (**Equation 4**).

### FRAP of Abl2 on MTs

Preparation of biotinylated MT seeds and functionalization of TIRF chambers were described as above. 10% biotinylated 10% rhodamine-labelled GMPCPP-stabilized seeds were flowed into the chamber and unattached seeds were washed out with BRB80. Imaging buffer (1 µM Abl2-eGFP, 1X anti-blink cocktail, 50 mM KCl, 0.02% methylcellulose, and BRB80) was perfused into the flow chamber and left to incubate at RT for 5 min to allow condensates to form on MTs. Prior to photobleaching, images were acquired at 1 fps for 15 seconds for both 488 and 561 nm channels. Photobleaching was carried out using 20% power of the 405 nm laser line, with 20 µs dwell time. Intensity of each ROIs were recorded at ∼0.37 fps over a 10 min acquisition period. Images were acquired using Nikon Elements software; and a Nikon Ti2-E inverted microscope equipped with a 100X 1.49NA TIRF objective, stage top Piezo for 405 nm FRAP laser, and a Photometrics Prime 95B CMOS camera. FRAP curves of MTs from 4 TIRF chambers were fitted to **Eqn. 4**.

### Microtubule nucleation observation using confocal microscopy

8 µM 10% Alexa Fluor 647-labeled porcine brain tubulin in BRB80 was incubated with the MT polymerization buffer (2 mM of GTP, 1 mM DTT, 10% glycerol), 1 µM Abl2-eGFP, 3% dextran at 37°C. Reactions were added to a 384-well glass-bottom plate (Corning, Corning, NY), which was acid-washed and coated with 2% F-127 for 30 min. Tubulin nucleation was monitored using Nikon CSU-W1 SoRa spinning disk confocal 30 min and 1 hr after the start of the reactions.

### Fixed-cell sample preparation

Cells were plated on 12 mm #1.5 coverslips (Warner Instruments, Hamden, CT) in 3 cm TC-treated dishes (Corning, Corning, NY). Coverslips were acid washed at 55°C overnight and coated with 50 µg*ml^-1^ poly-d-lysine for 20 min at room temperature and 10 µg*ml^-1^ fibronectin in phosphate-buffered saline (PBS) for 1 h at 37°C. Cells were treated with 10 µM nocodazole for 60 min, and the coverslips were taken out at 5 min after washout with complete medium. Cells were fixed with 4% paraformaldehyde in cytoskeleton buffer (10 mM MES pH 6.1, 138 mM KCl, 3 mM MgCl_2_, 2 mM EGTA, 0.32 M sucrose) for 20 min at room temperature and then permeabilized with permeabilization buffer (0.5% Triton X-100 in PBS) for 20 min. The fixed cells were blocked in blocking buffer (2% BSA, 0.1% Triton X-100 in PBS) followed by the primary and secondary antibodies. Primary a-tubulin antibody (DM1a) and GTP-tubulin antibody (MB11) were used at 1:500 dilution. Alexa647 goat anti-mouse secondary antibody from Invitrogen was used at 1:200 to label a-tubulin. All antibodies were diluted in blocking buffer.

### SEC-MALS analysis and Stokes radius analysis

To measure the particle size of purified Abl2 and Abl2-557-C in solution, we used an experimental setup including an upstream size exclusion chromatography (SEC, Superdex 200 increase 10/300 GL) coupled with a downstream multi-angle light scattering (MALS), which includes a multiple light scattering detector (DAWN HeleosII, Wyatt Technology Corporation) and a refractive index detector (Optilab T-rEX, Wyatt Technology). Each sample injection consisted of 0.5 to 2 mg of purified protein sample in buffer containing 20 mM HEPES pH 7.25, 5% glycerol, 100 mM KCl, 1 mM DTT. The flow rate was set at 0.4 mL/min. Data were recorded at 1 second interval and processed using ASTRA software (Wyatt Technology). Stokes radius analyses were performed as described previously^102^.

## Data analysis and availability

Fiji and MATLAB were used for all image analysis. At least triplicate technical replicates were performed for each experiment unless otherwise stated. Custom-written MATLAB scripts for lattice damage analysis are available upon request.

