## Supplemental Figures for "Abl2 mediates microtubule nucleation and repair via tubulin co-condensation"

**Figure S1. Characterization of the physical properties of Abl2 and interactions with tubulin. (A)** SEC analysis of standard proteins as indicated using the Superdex 200 increase 10/300 GL column (top). The standard linear plot of the standard proteins $log_{10}R_s$ and $K_{av}$ (bottom), with $R_s$ =Stokes radius and $K_{av} = (V_e-V_0)/(V_t-V_0)$, in which $V_e$ represents the elution volume of the protein peak, $V_0$ represents the column void volume (8 mL), and $V_t$ represents the total column volume (23.562 mL). Stokes radii of Abl2 (10.7 mL), Abl2-557-C (11.4 mL), and tubulin (14.2 mL) were calculated from the fitted eque. SDS-PAGE gels of the SEC samples are shown in the right panels. **(B, C)** SEC-MALS analysis of Abl2 in **(B)** and Abl2-557-C in **(C)** showed that Abl2 and Abl2-557-C mainly exist as monomers in solution despite the large Stokes radius. In **(B)**, Peak1 represents the predominant monomer peak; Peak2 represents a small amount of dimer formation; Peak3 represents Abl2 multimers. **(D, E)** All purified proteins used in this work were analyzed by SDS-PAGE and stained with Coomassie Blue G250 to assess purity.

**Figure S2. Abl2 promotes MT assembly and nucleation. (A)** At the termination of turbidity assays shown in Figure 3, they were subjected to ultracentrifugation. 1/100 of the resuspended volume was retrieved from the reactions to visualize the amount of pellet MTs in the reactions with or without Abl2. The intensity of the tubulin bands was quantified in the right panel. **(B)** WT and Abl2 KO COS-7 cells were lysed and 30 μg of the lysates were loaded for SDS-PAGE and western blot analysis. The left panel is Ponceau S staining showing the total protein loaded in each lane. The right panel is an immunoblot for Abl2 using the monoclonal Ar11 and shows the loss of Abl2 signal in Abl2 KO cells. **(C)** The average cell area of WT and Abl2 KO COS-7 does not differ significantly. The cell area was outlined on the intensity and the area was measured. n.s., no significance. **(D)** Representative images of FIJI analysis of MT recovery in cells are shown. A region of interest (ROI) was chosen in the cell body where no MT segments were observed, as shown by the red box in the left panels. Images were thresholded using 4X the background ROI intensity to pick up the saturated intensity that reflects recovered MTs, as shown by the red pixels in the right panels. The area of the red pixels was measured for each cell.

**Figure S3. MTs undergo more frequent rescue and repair in presence of Abl2**. **(A)** Shrinkage frequency (μm of shrinkage length per min) and rescue frequency (events per μm of shrinkage length) represented as line plots. Consistent with previous findings, Abl2 decreased depolymerization rate of MTs grown from 10.5 μM rhodamine tubulin relative to control ($p < 0.0001$). Whereas the tubulin-binding fragment 688-924-eGFP decreased and tubulin-binding deficient Abl2Δ688-790 did not impact rescue frequency relative to tubulin alone ($p < 0.0001$; n.s., respectively), tubulin- and MT-binding Abl2-eGFP increases rescue frequency by 2-fold relative to tubulin alone ($p < 0.0001$). From left to right: n = 91-315 MTs per sample. Means ± SEM shown as circular markers and error bars, respectively. **(B)** Distribution of healthy segment lengths of MTs stored at 23°C and 37°C overnight. Consistent with previous studies, structurally intact segments of longer lengths were observed from MTs stored at 37°C overnight. 37°C storage allows for residual tubulin to repair MT shafts whereas 23°C inhibits. Means of fitted gamma distributions shown as dashed lines (mean lengths of 3.34 μm for 23°C and 4.92 μm for 37°C-stored MTs). $n \geq 350$ segments per condition. **(C)** Mean global Abl2 fluorescence intensities on 23°C- and 37°C-stored MTs. Means shown as solid black horizontal lines. 25-75% quartiles shown as box plots. $n \geq 90$ filaments analyzed per condition. **(D)** Distributions of tubulin reporter (repair) lengths of 23°C- and 37°C-stored MTs in the absence and presence of 10.5 μM Abl2-eGFP. $n \geq 90$ filaments analyzed per condition**.** **(E)** Distributions of incorporation events per μm of 23°C- and 37°C-stored MTs in the absence and presence of 1 μM Abl2-eGFP. $n \geq 200$ repair events analyzed per condition.

#### Figure S1

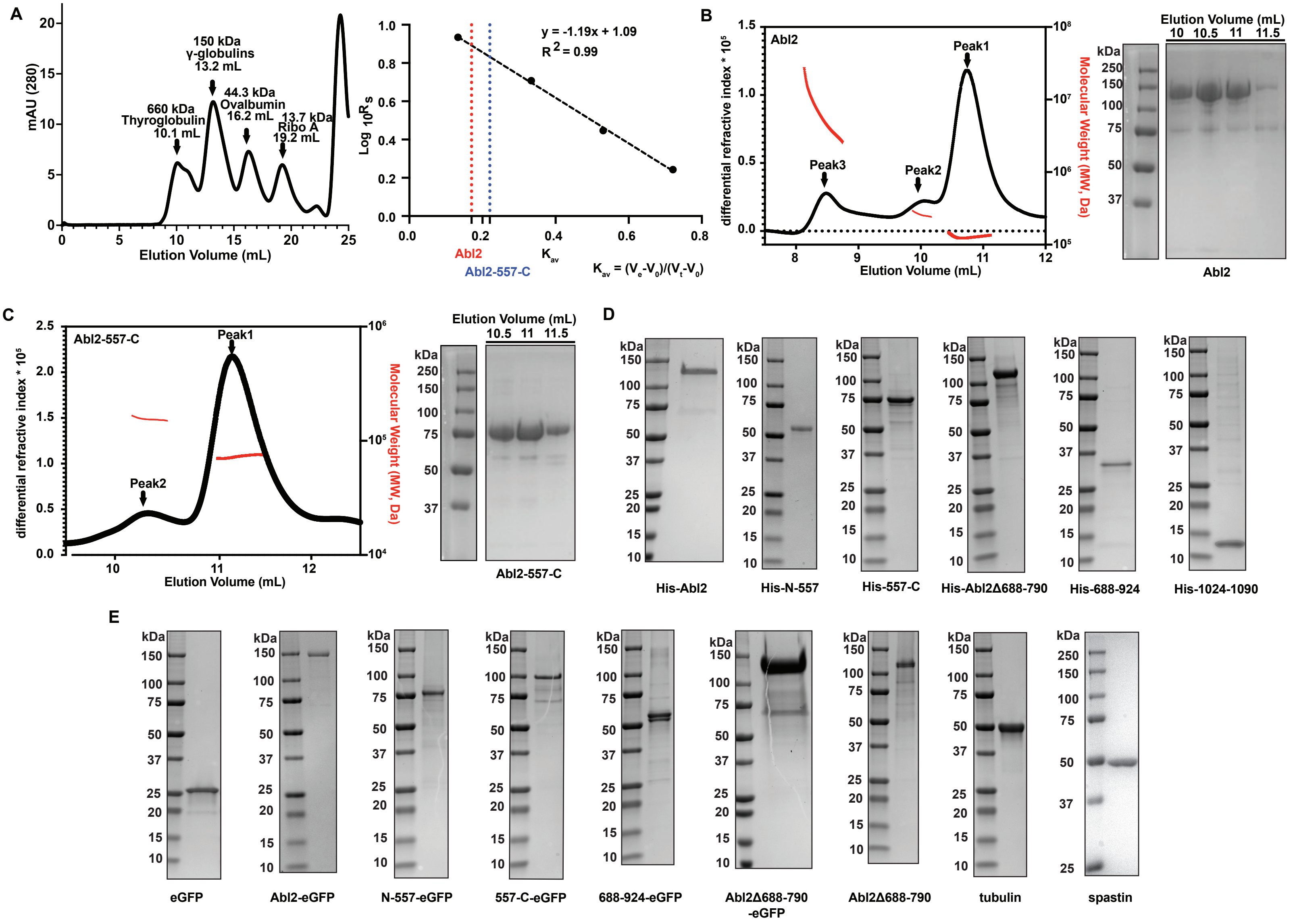
A
mAU (280)
Elution Volume (mL)
660 kDa
Thyroglobulin
10.1 mL
150 kDa
γ-globulins
13.2 mL
44.3 kDa
Ovalbumin
16.2 mL
13.7 kDa
Ribo A
19.2 mL
y = -1.19x + 1.09
R2 = 0.99
Abl2
Abl2-557-C
Kav
Kav = (Ve-V0)/(Vt-V0)
B
Abl2
differential refractive index * 10^5
Molecular Weight (MW, Da)
Peak1
Peak2
Peak3
Elution Volume (mL)
kDa
Abl2
C
Abl2-557-C
differential refractive index * 10^5
Molecular Weight (MW, Da)
Peak1
Peak2
Elution Volume (mL)
Abl2-557-C
D
His-Abl2
His-N-557
His-557-C
His-Abl2Δ688-790
His-688-924
His-1024-1090
E
eGFP
Abl2-eGFP
N-557-eGFP
557-C-eGFP
688-924-eGFP
Abl2Δ688-790
-eGFP
Abl2Δ688-790
tubulin
spastin

### Figure S2

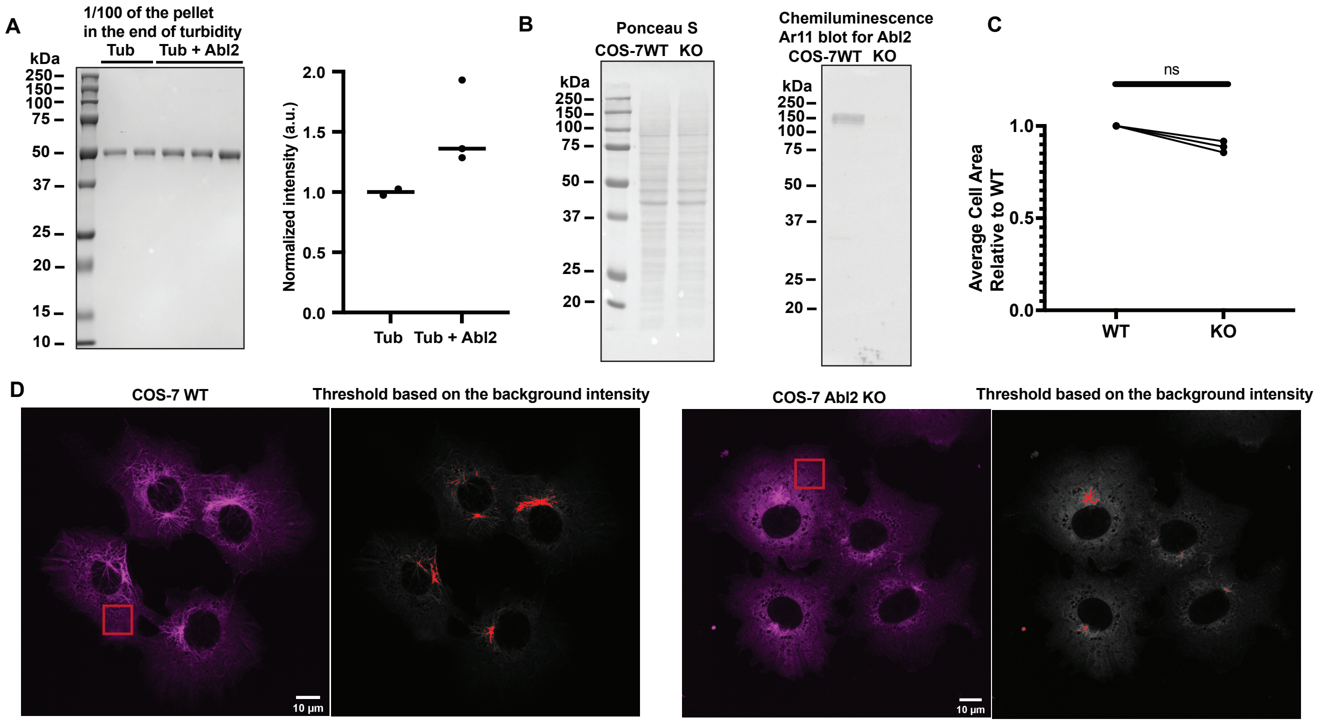
A
1/100 of the pellet
in the end of turbidity
Tub
Tub + Abl2
kDa
250
150
100
75
50
37
25
20
15
10
Normalized intensity (a.u.)
2.0
1.5
1.0
0.5
0.0
Tub
Tub + Abl2
B
Ponceau S
COS-7WT KO
kDa
250
150
100
75
50
37
25
20
Chemiluminescence
Ar11 blot for Abl2
COS-7WT KO
kDa
250
150
100
75
50
37
25
20
C
ns
Average Cell Area
Relative to WT
1.0
0.5
0.0
WT
KO
D
COS-7 WT
Threshold based on the background intensity
10 µm
COS-7 Abl2 KO
Threshold based on the background intensity
10 µm

### Figure S3

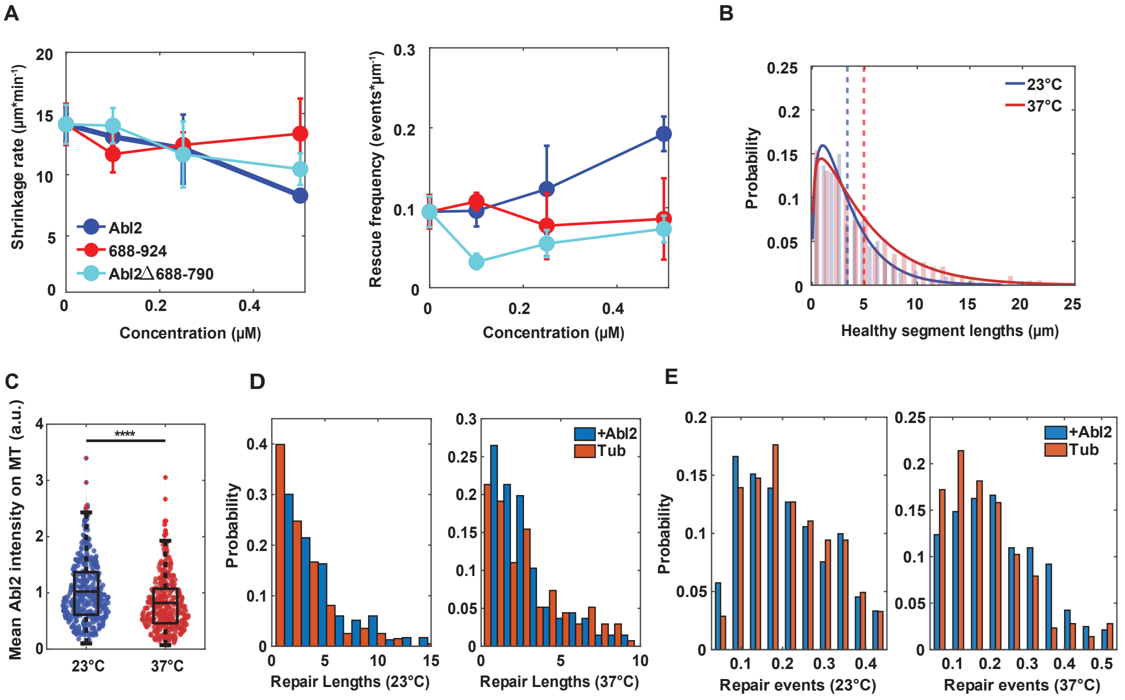
A
Shrinkage rate (µm*min-1)
Concentration (µM)
Abl2
688-924
Abl2△688-790
Rescue frequency (events*µm-1)
Concentration (µM)
B
Probability
Healthy segment lengths (µm)
23°C
37°C
C
Mean Abl2 intensity on MT (a.u.)
****
23°C
37°C
D
Probability
Repair Lengths (23°C)
Repair Lengths (37°C)
+Abl2
Tub
E
Probability
Repair events (23°C)
Repair events (37°C)
+Abl2
Tub
